# Genomic mechanisms and consequences of diverse postzygotic barriers between monkeyflower species

**DOI:** 10.1101/2023.04.21.537885

**Authors:** V. Alex Sotola, Colette S. Berg, Matt Samuli, Hongfei Chen, Samuel J. Mantel, Paul A. Beardsley, Yao-Wu Yuan, Andrea L. Sweigart, Lila Fishman

**Author notes:** Biology Department, State University of New York at Oneonta, Oneonta, NY 13820.

## Abstract

The evolution of genomic incompatibilities causing postzygotic barriers to hybridization is a key step in species divergence. Incompatibilities take two general forms – structural divergence between chromosomes leading to severe hybrid sterility in F_1_ hybrids and epistatic interactions between genes causing reduced fitness of hybrid gametes or zygotes (Dobzhansky-Muller incompatibilities). Despite substantial recent progress in understanding the molecular mechanisms and evolutionary origins of both types of incompatibility, how each behaves across multiple generations of hybridization remains relatively unexplored. Here, we use genetic mapping in F_2_ and RIL hybrid populations between the phenotypically divergent but naturally hybridizing monkeyflowers *Mimulus cardinalis* and *M. parishii* to characterize the genetic basis of hybrid incompatibility and examine its changing effects over multiple generations of experimental hybridization. In F_2_s, we found severe hybrid pollen inviability (< 50% reduction vs. parental genotypes) and pseudolinkage caused by a reciprocal translocation between Chromosomes 6 and 7 in the parental species. RILs retained excess heterozygosity around the translocation breakpoints, which caused substantial pollen inviability when interstitial crossovers had not created compatible heterokaryotypic configurations. Strong transmission ratio distortion and inter-chromosomal linkage disequilibrium in both F_2_s and RILs identified a novel two-locus genic incompatibility causing sex-independent gametophytic (haploid) lethality. The latter interaction eliminated three of the expected nine F_2_ genotypic classes via F_1_ gamete loss without detectable effects on the pollen number or viability of F_2_ double heterozygotes. Along with the mapping of numerous milder incompatibilities, these key findings illuminate the complex genetics of plant hybrid breakdown and are an important step toward understanding the genomic consequences of natural hybridization in this model system.

## Introduction

The evolution of postzygotic reproductive barriers, caused by incompatibilities between interacting genes and/or meiotic dysfunction in chromosomally divergent hybrids, is a key component of speciation (Coyne and Orr 2004; Fishman and Sweigart 2018; Coughlan and Matute 2020). Experimental hybridization of closely related species provides a window into the nature and origins of postzygotic barriers, revealing both the genetic mechanisms of genomic incompatibility and potential magnitude of barriers to gene flow upon secondary contact. The characterization of both early-and late-generation experimental hybrid populations is a particularly promising approach (Matute *et al*. 2020) Multi-generation approaches can capture extrinsic incompatibilities dependent on environmental context (Walter *et al*. 2020), replicate natural hybrid zones (Pritchard and Edmands 2012), and under controlled growth conditions, allow comparison of early and late hybrid generations. Additional rounds of recombination and distinct assemblages of hybrid genotypes in later generations provide increased power to detect and localize incompatibilities (Moyle and Nakazato 2008). Furthermore, comparing patterns of hybrid breakdown across experimental generations can reveal how initial selection against incompatible genotypes shapes hybrid genomes and whether genetic and chromosomal incompatibilities quickly resolve (e.g., sort to parental genotypes) after initial hybridization or are maintained as potentially costly polymorphisms.

Fitness breakdown in experimental hybrids can be characterized in two complementary ways: by direct measurement of fertility, viability, and other fitness metrics and by characterizing deviations from expected Mendelian allele or genotype frequencies (transmission ratio distortion; TRD). TRD is a common feature of experimental hybrid populations, and interspecific mapping populations often exhibit significant distortion across a large fraction of their chromosomes (Fishman and McIntosh 2019). In plants, TRD at individual loci has many documented sources, including environmental selection (Yin *et al*. 2004), meiotic drive by chromosomes (Fishman and Saunders 2008), exposure of suppressed gamete killers (Koide *et al*. 2008a), pollen competition (Fishman *et al*. 2008), and Dobzhansky-Muller incompatibilities that kill gametes or zygotes (Leppälä *et al*. 2008; Kerwin and Sweigart 2017). In some cases, strong Dobzhansky-Muller incompatibilities (e.g., gametic or gametophytic lethals) can generate both local distortion and linkage disequilibrium between physically unlinked loci if they kill a given multi-locus genotypic class (Colomé-Tatché and Johannes 2016). Although differentiating among multiple mechanisms of hybrid TRD can be challenging, joint mapping of fitness traits and TRD in multiple generations increases power and precision to characterize intrinsic postzygotic barriers.

Along with genic Dobzhansky-Muller incompatibilities, chromosomal rearrangements can play important direct and indirect roles in postzygotic breakdown. Inversions often do not cause direct underdominant effects on hybrid fertility in plants (Fishman and Sweigart 2018; Huang and Rieseberg 2020; Zhang *et al*. 2021), but their suppression of recombination across large genomic regions in hybrids can link multiple incompatibility loci and extend their barrier effects (Rieseberg 2001; Noor *et al*. 2001; Livingstone and Rieseberg 2004). Reciprocal translocations tend not to be as common as inversions within species, but often distinguish sister species of flowering plants (Grant 1971; Fishman *et al*. 2013; Ostevik *et al*. 2020). In contrast to inversions, translocations can directly cause meiotic dysfunction and sterility in F_1_ hybrids (Stathos and Fishman 2014); with or without crossovers, segregation of paired chromosomes with reciprocal translocations often produce at least 50% unbalanced gametes (Burnham 1956). Because novel translocations should be strongly disfavored by selection until reaching >50% frequency (Lande 1984), their initial establishment and spread in diverging populations has been a long-standing puzzle in evolutionary biology. However, some plant species maintain permanent translocation heterozygosity (Golczyk *et al*. 2014) and/or avoid adjacent segregation, so populations segregating for translocations may rapidly shift (genetically or epigenetically) to minimize the deleterious effects of heterokaryotypy. Thus, understanding how translocations contribute to post-zygotic breakdown and genomic transmission after initial hybridization (both directly and indirectly) is an important step in understanding the processes that maintain plant species barriers.

Here, we investigate the patterns and underlying mechanisms of hybrid breakdown in multiple generations of hybrids between closely-related monkeyflowers (Phin *Mimulus* section *Erythranthe*, *M. cardinalis* and *M. parishii* (also known as *Erythranthe cardinalis* and *E. parishii*)). Despite dramatic differences in floral morphology (Beardsley *et al*. 2003; Fishman *et al*. 2015; Liang *et al*. 2022), hummingbird-pollinated *M. cardinalis* and selfer *M. parishii* hybridize naturally and show signatures of recent introgression where their ranges overlap in southern California (Nelson *et al*. 2021b). The *Erythranthe* group, which also includes bee-pollinated *M. lewisii*, has been long-standing model for understanding speciation by pollinator shifts and other ecological factors (Hiesey *et al*. 1971; Bradshaw *et al*. 1995; Bradshaw and Schemske 2003; Angert *et al*. 2008) and has recently emerged as a model system for studying Asteridae floral development (Yuan *et al*. 2016; Yuan 2019; Liang *et al*. 2022). Because phylogenomic analyses reveal substantial genomic reticulation among all taxa despite rapid morphological and structural divergence (Nelson *et al*. 2021b), *Erythranthe* is a particularly rich system for investigating the origins, maintenance, and interactions of both pre-and postzygotic species barriers.

As a platform for investigations of both postzygotic barriers (this study) and dramatic floral, life history, and vegetative divergence, we generate dense genetic linkage maps of *M. parishii* x *M. cardinalis* F_2_ and recombinant inbred line (RIL) hybrids using genome-wide ddradseq anchored in a new *M. cardinalis* genome assembly (mimubase.org). Previous coarse mapping in hybrids of each focal species with *M. lewisii* (Fishman *et al*. 2013, 2015), plus genotyping of breakpoint markers in a small number of *M parishii* x *M. cardinalis* hybrids (Stathos and Fishman 2014) suggest that these naturally hybridizing taxa are distinguished by at least one reciprocal translocation. High-resolution genome-wide mapping of male fertility (pollen viability and number) and TRD in each generation allows us to address specific hypotheses about putative barriers, identify the full complement of genic and chromosomal incompatibilities genome-wide, and track their evolution and influence from F_2_ to advanced generation hybrids.

Specifically, we ask: Are patterns of postzygotic breakdown (hybrid sterility and transmission ratio distortion) consistent in multiple environments (i.e., between differnt F_2_ populations) and through multiple generations of self-fertilization (F_2_ hybrids vs. RILs)? Do chromosomes with reciprocal translocation sort to parental or other meiotically compatible karyotypes in advanced-generation hybrids, or does chromosomal underdominance persist? Are major incompatibility loci shared with other *Mimulus* section *Erythranthe* hybrids, or are there novel postmating reproductive barriers unique to this florally-divergent cross?

## Methods

### Study system and plant lines

The monkeyflowers of the *M. cardinalis* species complex (*Mimulus* section *Erythranthe;* Phrymaceae) are a well-established model system for understanding the genetics of floral evolution and speciation (Yuan 2019). Hummingbird-pollinated *M. cardinalis* has a broad latitudinal range in the western United States with distinct Arizonan, Southern Californian, and Sierran clades (Nelson *et al*. 2021b)*. Mimulus cardinalis* is parapatric in the Sierras with bee-pollinated high-elevation specialist *M. lewisii*, where reproductive isolation is maintained by habitat and pollinator preferences (Hiesey *et al*. 1971; Ramsey *et al*. 2003; Bradshaw and Schemske 2003), as well as two reciprocal translocations causing underdominant F_1_ hybrid sterility (Fishman *et al*. 2013; Stathos and Fishman 2014). *Mimulus parishii* is a small-flowered annual selfer restricted to Southern California, where it co-occurs with *M. cardinalis* in ephemerally wet desert washes. Hybrids between *M. cardinalis* and *M. parishii* have been observed in the field, and local sharing of identical organellar genomes (as well as nuclear reticulation) indicates an extensive history of mating and introgression despite extreme differences in floral morphology (Nelson *et al*. 2021b).

The plants in this study were all derived from two highly (>10 generations) inbred lines of Sierran *M. cardinalis* (CE10) and *M. parishii* (PAR), which were also used in previous investigations of species barriers (Bradshaw *et al*. 1998; Schemske and Bradshaw 1999; Ramsey *et al*. 2003; Bradshaw and Schemske 2003; Fishman *et al*. 2013, 2015; Nelson *et al*. 2021a). We generated PAR x CE10 F_1_ hybrids by hand-pollination (with prior emasculation of the PAR seed parent in the bud) and F_2_ hybrids by self-pollination of F_1_ hybrids. The F_2_ hybrids were grown in two separate greenhouse common gardens at the University of Montana (UM-F_2_; total *N* = 524) and the University of Connecticut (UC-F_2_ *N* = 253), along with parental control lines, and were phenotyped for numerous floral and vegetative traits including the pollen fertility traits presented here. Recombinant inbred lines (RILs) were generated by single-seed-descent from additional F_2_ individuals grown at the University of Georgia and California State Polytechnic University, Pomona; a total of 167 RILs were formed through 3-6 generations of self-fertilization.

### DNA extraction and sequencing

Genomic DNA was extracted from bud and leaf tissue of the greenhouse-grown F_2_ and RIL mapping populations using a CTAB chloroform protocol modified for 96-well plates (dx.doi.org/10.17504/protocols.io.bgv6jw9e). We used a double-digest restriction-site associated DNA sequencing (ddRADSeq) protocol to generate genome-wide sequence clusters (tags), following the BestRAD library preparation protocol (dx.doi.org/10.17504/protocols.io.6awhafe), using restriction enzymes PstI and BfaI (New England Biolabs, Ipswich, MA). Post-digestion, half plates of individual DNAs were labeled by ligation of 48 unique in-line barcoded adapters, then pooled for size selection. Libraries were prepared using NEBNext Ultra II library preparation kits for Illumina (New England BioLabs, Ipswich, MA). Each pool was indexed with a unique NEBNext i7 adapter and an i5 adapter containing a degenerate barcode and PCR amplified with 12 cycles. The F_2_ libraries were size-selected to 200-700bp using BluePippin 2% agarose cassettes (Sage Science, Beverly, MA) and sequenced (150-bp paired-end reads) in a partial lane of an Illumina HiSeq4000 sequencer at GC3F, the University of Oregon Genomics Core Facility. The RIL library was sequenced (150-bp paired-ends) without size-selection on an Illumina HiSeq4000 at Genewiz (South Plainfield, NJ).

### Sequence processing and linkage mapping

After sequencing, two separate ddRAD datasets were analyzed: one with samples from both F_2_ populations (*N* = 283 UM-F_2_ hybrids with 3 *M. parishii* and 2 *M. cardinalis* controls, and 253 UC-F_2_ hybrids, with 3 each F_1_, *M. parishii* and *M. cardinalis* controls) and one with samples from the RIL population (*N* = 167). Samples from both datasets were demultiplexed using a custom Python script (dx.doi.org/10.17504/protocols.io.bjnbkman), trimmed using Trimmomatic (Bolger *et al*. 2014), mapped to the *M. cardinalis* CE10 v2.0 reference genome (http://mimubase.org/FTP/Genomes/CE10g_v2.0) using BWA MEM, and indexed using SAMtools (Li *et al*. 2009). The RIL dataset was also filtered in SAMtools using a mapping quality ≥ 29. We called SNPs in both datasets using HaplotypeCaller in GATK v3.3 in F_2_s, v4.1.8.1 in RILs (McKenna *et al*. 2010).

Next, we performed a series of filtering steps to generate sets of high-quality SNPs. In the F_2_ dataset, we filtered using vcftools (Danecek *et al*. 2011), retaining sites with read depth ≥ 5, mapping quality ≥ 10, and < 40% missing data. We also filtered out loci deviating from Hardy-Weinberg Equilibrium at a p-value < 0.00005. In the RIL dataset, we filtered a combined GVCF file using GATK, retaining sites with read depth ≥ 4\**N* (with *N* = number of RIL samples) and < the mean + 2 * the standard deviation, QD score < 2.0, FS score > 60, MQ < 40, MQ rank sum <-12.5, ReadPosRankSum <-8.0, and < 10% missing genotypes. For both datasets, we used custom scripts to remove sites that were not polymorphic in the parents and heterozygous in the F_1_ hybrids (F_2_: https://github.com/bergcolette/F2_genotype_processing). We excluded individuals from the F_2_ dataset with > 10% missing data and from the RIL dataset with low coverage, high missingness, or excessive heterozygosity (> 50%, indicating line contamination). These filtering steps produced an F_2_ dataset with 18,119 SNPs (*N* = 252 UM-F_2_ and 253 UC-F_2_) and a RIL dataset with 47,851 SNPs (*N* = 145).

To produce sets of high-quality marker genotypes for mapping, we binned each dataset into 18-SNP windows using custom Python and R scripts (provided at github links above), requiring ≥ 8 sites to have SNP genotype calls to assign a windowed genotype. In the F_2_ binning script, *M. cardinalis* homozygotes were coded as 2, *M. parishii* homozygotes as 0, and heterozygotes as 1. We called windows with mean values < 0.2 as *parishii* homozygotes, > 1.8 as *cardinalis* homozygotes, and between 0.8 and 1.2 as heterozygotes. Windows with means outside of these ranges were coded as missing genotypes. For the RILs, we required ≥ 88% of SNP calls to match each other to assign each parental homozygous genotype (*e.g*., 16/18 sites = homozygous for *M. cardinalis* alleles; https://github.com/vasotola/GenomicsScripts).

We generated linkage maps for each dataset using Lep-MAP3 (Rastas 2017). First, we used the *SeparateChromosomes2* module to assign markers to linkage groups (F_2_: LodLimit = 25, theta = 3, RIL: LodLimit = 28, theta = 0.2). In the RIL dataset, 10 markers were assigned to linkage groups inconsistent with the reference genome assembly; we manually re-assigned these markers to linkage groups corresponding to their reference assembly chromosomes. Next, we performed iterative ordering using the *OrderMarkers2* module (Kosambi mapping function; 6 iterations/per linkage group in the F_2_s, 10 in the RILs); the order with the highest likelihood for each linkage group was chosen. This resulted in an F_2_ map with 997 markers in seven linkage groups, and a RIL map with 2,535 markers in eight linkage groups. In the RIL dataset, the genotype matrix output by Lep-MAP3 differed in two important respects from the input file. First, due to stringent thresholds for calling windowed genotypes, our input file includes a high percentage of missing data (23% of genotypes are coded as ‘no call’), whereas the output file contains no missing data (Lep-MAP3 converts each ‘no call’ genotype to a called genotype). Second, the Lep-MAP3 output file contains more heterozygous genotype calls than the input file. The reason for this increase in heterozygosity is that Lep-MAP3 disproportionately converts ‘no call’ genotypes to heterozygotes: relative to the input file, the output genotype matrix includes 115% more heterozygotes, compared to only 18% more *M. cardinalis* homozygotes and 20% more *M. parishii* homozygotes. Notably, Lep-MAP3 frequently converted ‘no call’ genotypes to heterozygotes when they occur at single markers between recombination breakpoints. Because most recombinational switches in this RIL population are between alternative homozygotes, any window that contains an actual breakpoint will carry a mixture of *M. cardinalis* and *M. parishii* homozygotes at the 18 SNPs (and thus be coded as ‘no call’ in our windowed genotype matrix). To circumvent these problems, for all downstream analyses, we used a modified version of the genotype matrix output from Lep-MAP3 in which genotypes were recoded as ‘no call’ as in the input file.

### Transmission ratio distortion and linkage disequilibrium

Genotype frequencies were calculated, plotted, and tested for significant deviation from Mendelian expectations (*X*^2^ with 2 df) separately for the two F_2_ populations. TRD due solely to extrinsic factors (e.g., selection on parental alleles affecting germination under different conditions) may be distinct between the F_2_ grow-outs (Fishman and McIntosh 2019), whereas TRD due to intrinsic incompatibilities should be shared. For the RILs, we conducted parallel tests with expectations for alternate homozygotes set to 0.47 and heterozygotes to 0.06, which is the expectation given our final composition of RILs (nine individuals with three generations of selfing, 116 with four, 19 with five, and one individual with six generations of selfing). For all sets, we used both uncorrected (a = 0.01; critical value = 6.635) and stringent (Bonferonni-corrected; F_2_ critical value = 16.442, RIL critical value = 18.217) chi-squared tests to assess the significance of distortion.

We used the package *pegas* (Paradis 2010) in Program R (Team 2021) calculate pairwise linkage disequilibrium (*r*) between all markers within the UM-F_2_ and RIL mapping populations. To minimize false positives driven by the large number of pairwise comparisons, we plotted only those *r* values greater than the 95% quantile after bootstrapping 1000 times. All plots were visualized with R.

### QTL mapping of pollen traits

In the UM-F_2_ and RIL populations, we directly assessed male fertility by collecting all four anthers of the first flower from each plant into 50 ml of lactophenol-aniline blue dye. We counted viable (darkly stained) and inviable (unstained) pollen grains using a hemocytometer (≥ 100 grains/flower). We estimated total pollen grains per flower (count per ml x 50) and pollen viability as viable grains/total counted. For a handful of individuals with < 100 pollen grains counted, pollen viability was scored as missing data. Phenotyping in the RIL population was performed on siblings or selfed progeny of the individuals genotyped. In a few cases in the RILs (*N* = 12), the first flower had < 100 pollen grains, so both traits were instead collected from the second flower.

We mapped pollen QTLs in Windows QTL Cartographer 2.5 (Wang et al. 2012) using composite interval mapping (CIM; Zeng 1993, 1994), with forward-backward stepwise regression, a window size of 10 cM, five background markers, and a 1-cM walk speed. We used permutations (*N* = 1000) to set genome-wide significance thresholds for QTL peaks and calculated 1.5-LOD-drops to determine confidence intervals for QTL locations. Because pollen viability QTLs exhibited complex interactions in F_2_s (see Results), we directly estimated QTL effects (at each peak marker) and interactions using the Generalized Linear Model module in JMP16 (SAS Institute, Cary, NC), starting with a full factorial model of all QTLs and 2-way interactions and removing nonsignificant (P > 0.05) interactions. To test for pollen viability signatures of a two-locus gametic incompatibility (LG4-LG8) detected from TRD and LD patterns, we contrasted Least Squared Means for each extant F_2_ class from an ANOVA also including the large-effect translocation breakpoint marker at 60.55Mb on Chr 6.

QTL mapping of underdominant effects in RILs has low power due to (relatively) few heterozygotes, plus most RIL QTL-mapping algorithms (including those in WinQTLCart) exclude non-homozygous genotypes. To directly test for persistent underdominant effects of the Chr 6-7 translocation in RILs, we screened for genotype-pollen viability associations at uniquely positioned markers with a joint sample size >70 across the breakpoint region (57.07-61.08 Mb of Chr6 and 7.51-13.05 of Chr 7; n = 42), using t-tests in the response screening module in JMP16 (SAS Institute, Cary, NC). However, we caution that because RIL hybrid male sterility was measured on siblings or descendants of the genotyped individuals, underdominant effects will likely be underestimated. That is, regions genotyped as heterozygous might actually be homozygous in phenotyped individuals, potentially explaining why a few of them are highly fertile. We conservatively controlled for multiple tests by using an FDR-corrected *P*-value of 0.05.

## Results

### Comparative linkage mapping

Across most of the genome, the F_2_ and RIL genetic maps are highly collinear, with linkage groups and marker order largely reconstructing the physical order of the eight chromosomes of the *M. cardinalis* reference genome (Figures S1& S2). The notable exception to this pattern involves markers on Chromosomes (Chr) 6 and 7, where previous work suggested an *M. cardinalis*-specific reciprocal translocation vs. both *M. parishii* and *M. lewisii* (Fishman *et al*. 2013, 2015; Stathos and Fishman 2014). There is no single linear order of markers in heterozygotes for a reciprocal translocation (Livingstone *et al*. 2000), which generates linkage across the entire involved chromosomes during F_1_ meiosis and a continuous LG 6&7 in the F_2_ hybrids. In the RILs, the portion of Chr 6 distal to the translocation forms a distinct linkage group, while tight linkage between markers on the end of Chr 6 and the first 8 Mb of Chr 7 in *M. cardinalis* (purple and green/red segments, respectively, in Figure 1) generates a second composite linkage group. The RIL map is ∼35% longer than the F_2_ map (total length = 892.95 cM vs. 661.37 cM), consistent with the additional generations of recombination. In both maps, recombination rates per Mb are dramatically lower (near zero) across the central 20-40 Mb of each chromosome (Figure S1). This pattern is also evident for all chromosomes other than Chr 7 in an intraspecific *M. cardinalis* map (Nelson *et al*. 2021b; a), consistent with a primarily metacentric chromosomal structure in *Erythranthe* species.

**Figure 1.**
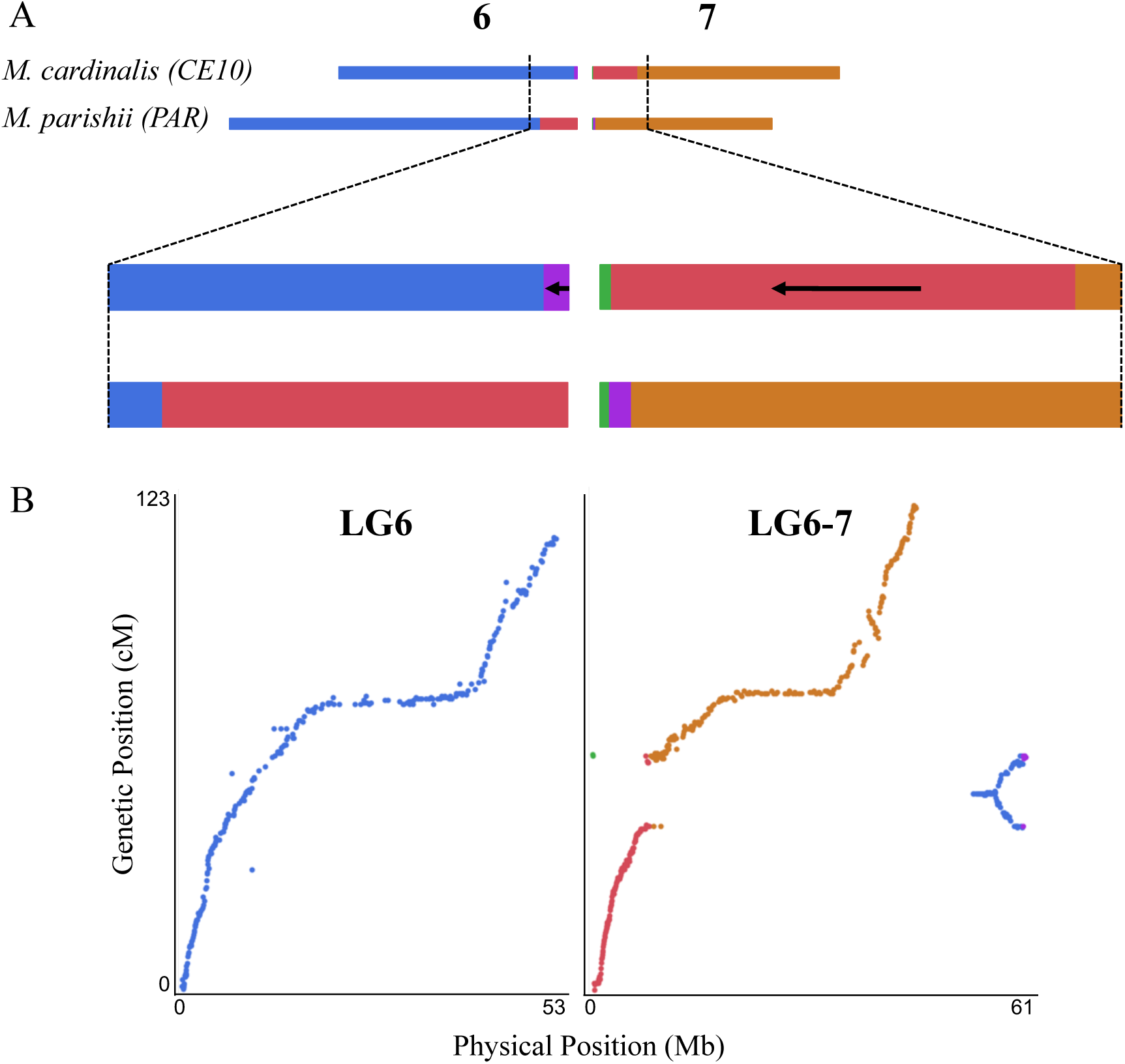
*Mimulus cardinalis* (CE10) and *M. parishii* (PAR) exhibit a reciprocal translocation between Chromosomes (Chr) 6 and 7. *A*) Schematic showing the affected chromosomes and regions: the distal region of Chr 6 in PAR has moved to the proximal region of 7 in CE10 (red) and a smaller proximal region of Chr 7 in PAR has moved to the distal-most region of Chr 6 in CE10 (purple). Black arrows indicate blocks where the CE10 order is inverted relative to the ancestral PAR order. Genome coordinates for the colored blocks are given in Table S1. *B*) Plot of physical position (Mb, CE10g_v2.0; www.Mimubase.org) by genetic position for RIL linkage groups 6 and 6-7. SNP markers (dots) are colored by the locations defined in *A*. RIL LG6 includes only markers from the collinear region of Chr 6. RIL LG6-7 includes all markers from the translocated regions, but the genetic order reflects multiple distinct linkages associated with the divergent parental chromosomes (e.g., blue markers near the translocation breakpoint at ∼60 Mb in CE10 are tightly linked to markers of all other colors).

### Genome-wide patterns of transmission ratio distortion (TRD)

All linkage groups, with the exception of LG2 in the UM-F_2_ mapping population, exhibited strong transmission ratio distortion (TRD) in all three maps (Figure 2, Figure S3, Tables S2-S3). Patterns of TRD were very similar between the two F_2_ populations, suggesting that TRD is largely caused by intrinsic factors (e.g., incompatibilities) vs. environmental selection (e.g., germination conditions). The two F_2_ grow-outs shared regions of excess *M. cardinalis* homozygosity on LGs 1, 4, 5 and 6 (distal to translocation), excess *M. parishii* on LG8, and excess heterozygosity on LG3 and in the translocation region on LG6&7. The UC-F_2_ also exhibited excess heterozygosity and *M. cardinalis* alleles on LG2. Additional generations of recombination and selection in the RIL population strengthened and refined the locations of shared TRD peaks on LGs 1, 2, 3, 6&7, and 8. However, RILs exhibited a novel region of very strong *M. parishii* excess on one arm of LG5; this signal was absent or opposite in the F_2_ hybrids, suggesting an advantage of *M. parishii* alleles in this region specific to either the environment or the increasingly homozygous genetic background of RIL formation. In addition to local peaks of TRD, the RIL population exhibited slightly higher residual heterozygosity across the genome (9.3%) than the Mendelian expectation (6%) (Figure 2). While some of this overall excess is due to major local peaks in heterozygosity shared with the F_2_ hybrids (e.g., on LG3 and in the LG6&7 breakpoint region, Figure 2), the remainder may reflect genome-wide selection against weakly incompatible parental homozygous combinations in the RILs (Thompson *et al*. 2022).

**Figure 2.**
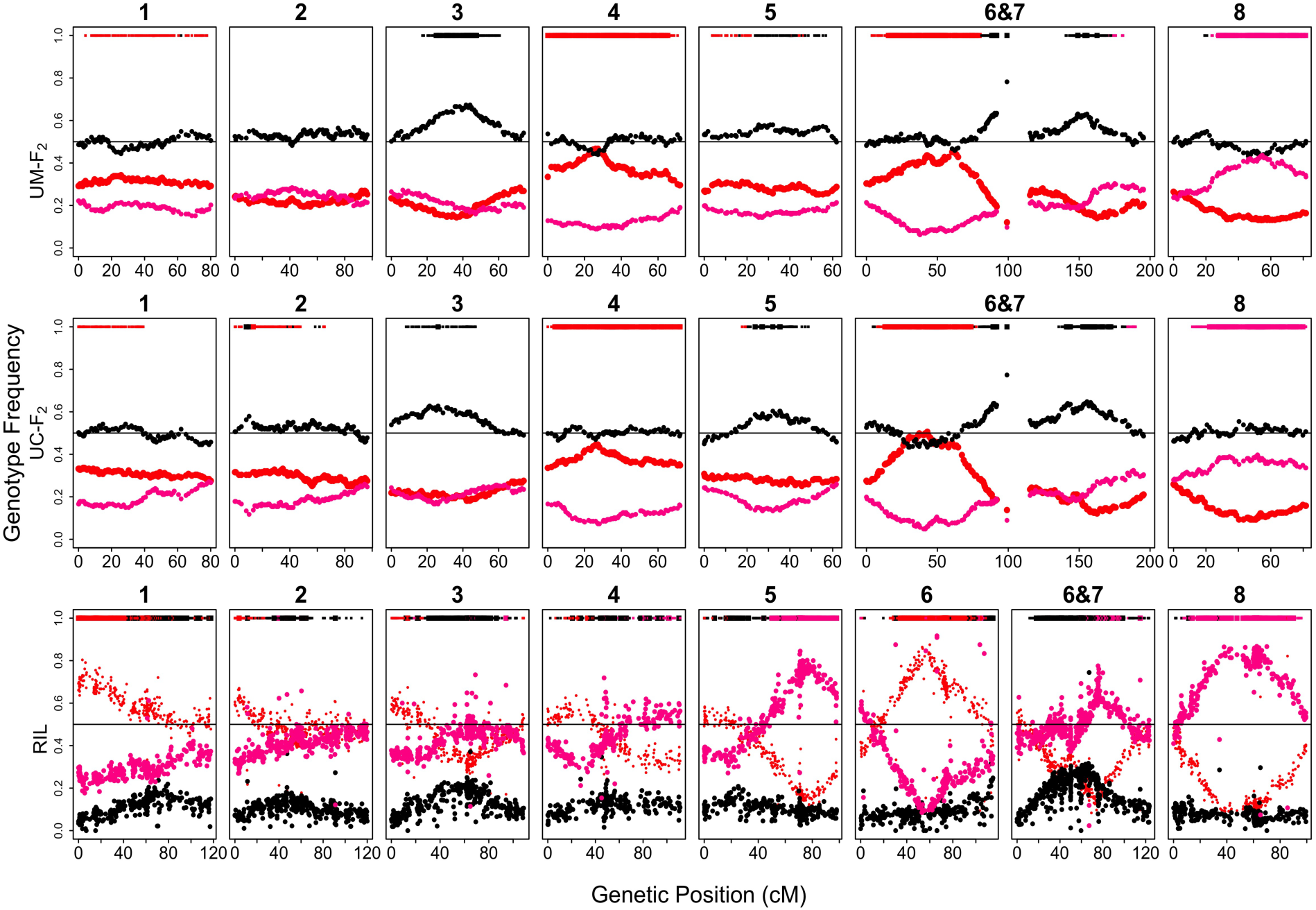
Genotype frequencies in the F_2_ and RIL populations across the genome. Red dots indicate homozygous *M. cardinalis*, pink is homozygous *M. parishii*, and black is heterozygous. The bars at the top of each plot indicate regions with significant transmission ratio distortion by χ^2^ tests at a = 0.01 (thin lines, critical value = 6.635) and at a more stringent, Bonferonni-corrected level (thick lines, F_2_ critical value = 16.442, RIL critical value = 18.217). Bar color indicates an excess of *M. cardinalis* homozygotes (red), *M. parishii* homozygotes (pink), or heterozygotes (black).

### Tests of hybrid incompatibilities as the source of TRD

To investigate whether peaks of TRD were caused by major two-locus hybrid incompatibilities (e.g. gametic or zygotic lethals), we characterized pairwise linkage disequilibrium (LD) between all markers in the UM-F_2_ and RIL populations (Figure 3, Tables S4 and S5). In the F_2_ hybrids, there was significant off-diagonal LD over most of LG6 and 7 (due to the translocation), between LG4 and the center of LG8, and between the distal ends of LG2 and LG8. In the RILs, parallel TRD on different chromosomes (e.g., near fixation of *M. parishii* alleles on both LG5 and LG8) can cause high inter-chromosome LD even without epistatic selection during RIL formation. Thus, it is not surprising that, in addition to sharing the F_2_ LD regions, the RILs exhibit more abundant and diffuse LD genome-wide (Figure 2, Table S5). However, some of these regions may house multi-locus incompatibilities that are too mild and/or late-acting to be statistically detected in the F_2_ generation (e.g., F_2_ sporophytic hybrid sterility)

**Figure 3.**
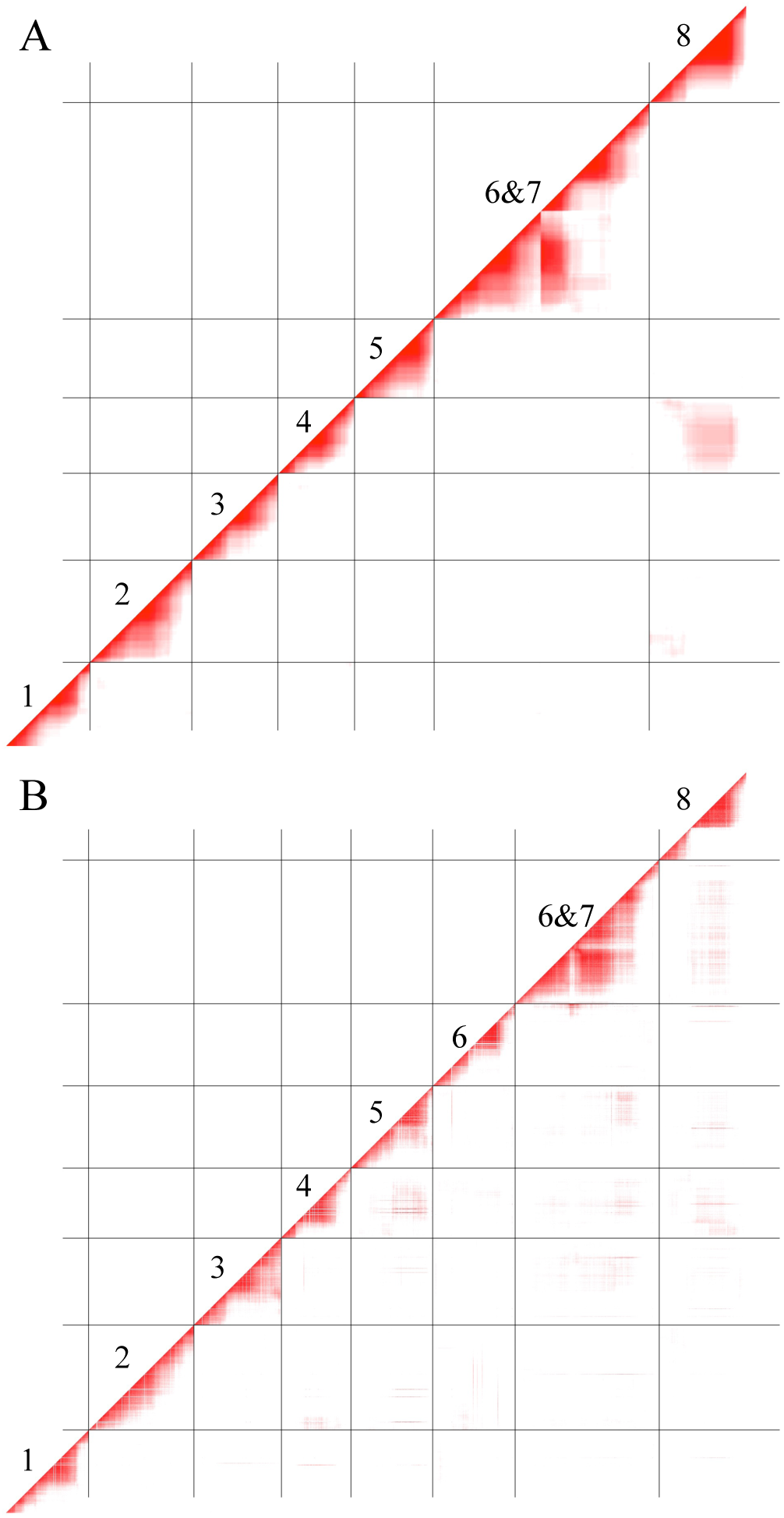
Linkage disequilibrium heatmaps for *M. parishii* x *M. cardinalis* UM-F_2_ (A) and RIL (B) populations, ordered by linkage group. Data shown are *r* values greater than the 95% quantile after bootstrapping (95% F2: 0.176; 95% RIL: 0.305). Values range from 0 (white) to 1 (bright red).

Strong and opposite TRD on LG4 (excess *M. cardinalis*) and LG8 (excess *M. parishii*) in the F_2_ hybrids (Figure 2), as well as high and persistent LD between these regions (Figure 3), suggested a major two-locus incompatibility. To investigate further, we tallied all two-locus genotype combinations at TRD-peak markers (6.8 Mb on Chr 4, 40 Mb on Chr 8; Figure 2, Figure S3) in both the UM-F_2_ and RIL populations (Table 1). These regions clearly deviate from the Mendelian expectation of independent segregation, with three of the expected nine genotypic classes entirely absent from both populations (Table 1). To test for the mechanism, we calculated the expected genotype frequencies under two different scenarios: 1) a gametic incompatibility in which all haploid gametes (both pollen and ovules) are inviable if they carry the *M. parishii-M. cardinalis* allelic combination at LG4 and LG8 (i.e., P;C gametes missing), and 2) a two-locus dominant zygotic incompatibility in which hybrids die if they carry at least one *M. parishii* allele at LG4 and at least one *M. cardinalis* allele at LG8 (P_; C_ zygotes die). In the F_2_ population, genotype frequencies best fit the sex-independent gametic incompatibility model, which distinctively predicts the numerous double heterozygotes observed (Table 1). The RIL genotypes are also most consistent with a sex-independent gametophytic mechanism; however, the observed excess of *M. parishii* transmission on LG8 exceeds that expected from a gametic incompatibility alone.

**Table 1.** Two-locus segregation pattern for loci on LG4 (6.75-7.65 Mb) and LG8 (12.87-40.08 Mb) showing opposite single-locus transmission ratio distortion and strong LD in *M. parishii* x *M. cardinalis* F_2_ hybrids and RILs. For each population, observed counts of individuals with each of the nine possible genotypes (C = *M. cardinalis* homozygote, P = *M. parishii* homozygote, and H = heterozygote) are compared to expected counts under normal Mendelian inheritance (Mendelian), expected counts under a model of sex-independent gametic inviability (P;C gametes inviable), and expected counts under a model of dominant zygote inviability (P_;C_ zygotes inviable). Counts are shown rounded to the nearest integer. See Supplemental Table S6 for unrounded counts and frequencies.

| Genotype<br>(LG4;LG8) | UM-F<sub>2</sub> Population |  |  |  | RIL Population |  |  |  |
| --- | --- | --- | --- | --- | --- | --- | --- | --- |
|  | Observed | Mendelian | P;C<br>gametes<br>inviable | P_<sub>;</sub>C_<sub>;</sub><br>zygotes<br>inviable | Observed | Mendelian | P;C<br>gametes<br>inviable | P_<sub>;</sub>C_<sub>;</sub><br>zygotes<br>inviable |
| C;C | 24 | 16 | 29 | 37 | 18 | 31 | 42 | 42 |
| H;C | 0 | 32 | 0 | 0 | 0 | 2 | 0 | 0 |
| P;C | 0 | 16 | 0 | 0 | 0 | 31 | 0 | 0 |
| C;H | 61 | 32 | 57 | 73 | 6 | 2 | 4 | 6 |
| H;H | 53 | 64 | 57 | 0 | 6 | 4 | 4 | 0 |
| P;H | 0 | 32 | 0 | 0 | 0 | 2 | 0 | 0 |
| C;P | 27 | 16 | 29 | 37 | 40 | 31 | 42 | 42 |
| H;P | 58 | 32 | 57 | 73 | 21 | 2 | 4 | 6 |
| P;P | 34 | 16 | 29 | 37 | 47 | 31 | 42 | 42 |

### Genetics of male fertility traits in hybrids

F_2_ hybrids (UM-F_2_) exhibited substantial variation for pollen viability (mean = 0.48, but significantly non-normal by Shapiro-Wilks test, P < 0.0001; Figure S4), whereas both parental lines were relatively fertile (mean = 0.73 for CE10 and 0.84 for PAR, *N* = 2 and 8, respectively). A major underdominant pollen viability QTL mapped to the translocation breakpoints on LG6&7, along with smaller QTLs on LGs 2, 3, and 4 (Figure 4; Table 2). To investigate the contribution of epistasis to the observed pollen inviability and test for unlinked modifiers of the translocation-associated underdominant sterility, we conducted a model selection analysis starting with a factorial model of the four peak markers and all two-way interactions. Only interactions including LG6&7 were significant at a = 0.05 (along with all four single QTLs), so we interpret QTL effects under that reduced model (*r^2^* = 0.42). The LG6&7 QTL was strongly underdominant (Table 2), while the other three pollen viability loci exhibited at least partial dominance of the *M. parishii* allele, with *M. cardinalis* homozygotes significantly more fertile than heterozygotes and *M. parishii* on LG4, significantly less fertile on LG3, and heterozygotes most fertile (but not significantly different from *M. cardinalis*) on LG2. The interactions involving the QTL on LG6&7 did not appear to modify the low fertility of translocation heterozygotes, which was consistent across genetic backgrounds (i.e., LG6&7 heterozygotes with alternative genotypes elsewhere were indistinguishable by Tukey’s Honest Significant Difference tests; Figure S5). Instead, the interactions involved idiosyncratic combinations of other genotypes, generally with low sample sizes; for example, F_2_ hybrids with *M. parishii* genotype at QTL PV3 and *M. cardinalis* at PV6&7 were about twice as fertile (Least Squared Mean [LSM] = 0.86 ± 0.11, *N* = 2) as those with the opposite homozygous combination (LSM = 0.45 ± 0.06, *N* = 8).

**Figure 4.**
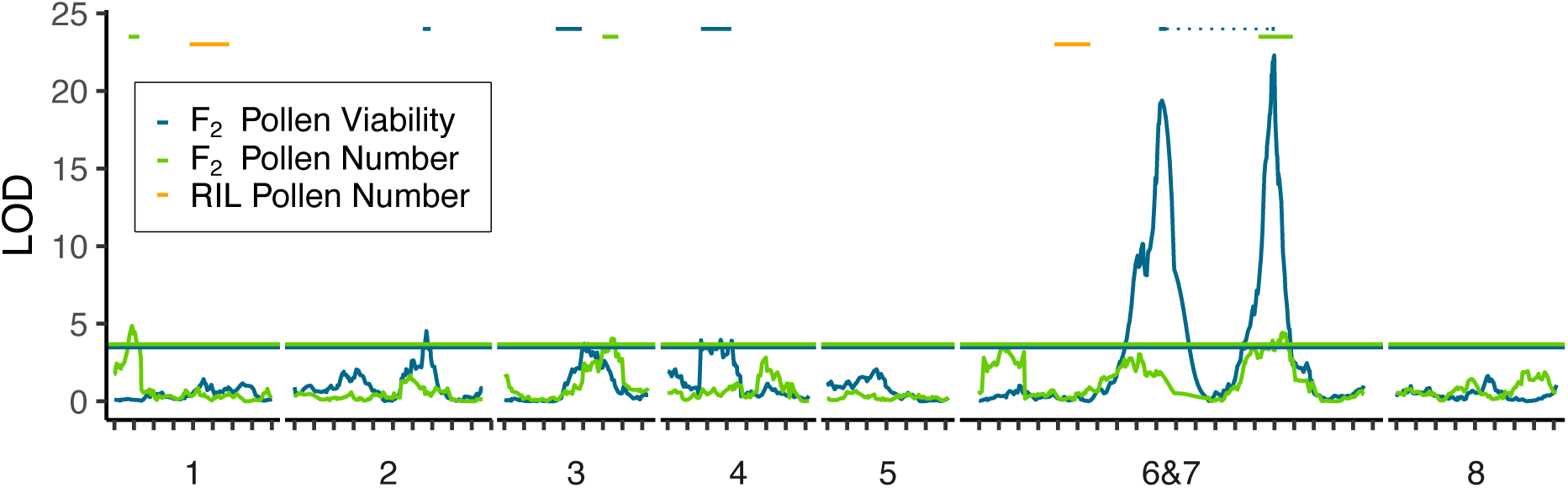
Quantitative trait loci (QTLs) for pollen viability (blue) and pollen number (green) in *M. parishii* x *M. cardinalis* F_2_ hybrids, as well as the locations of two pollen number QTLs in descendant RILs (orange, aligned by the physical position of markers). Horizontal lines denote the permutation-derived (N = 1000) LOD significance thresholds for each F_2_ trait, and the bars indicate QTL confidence intervals (1.5 LOD drop). The single major underdominant pollen viability QTL associated with the LG6&7 translocation maps to markers near both breakpoints (QTL locations connected by light dotted line), which cannot be linearly resolved in F_2_ hybrids. Because the RIL QTL mapping algorithm excluded heterozygotes, no RIL pollen viability QTLs were detected despite the negative effects of retained heterozygosity (see text).

**Table 2.**
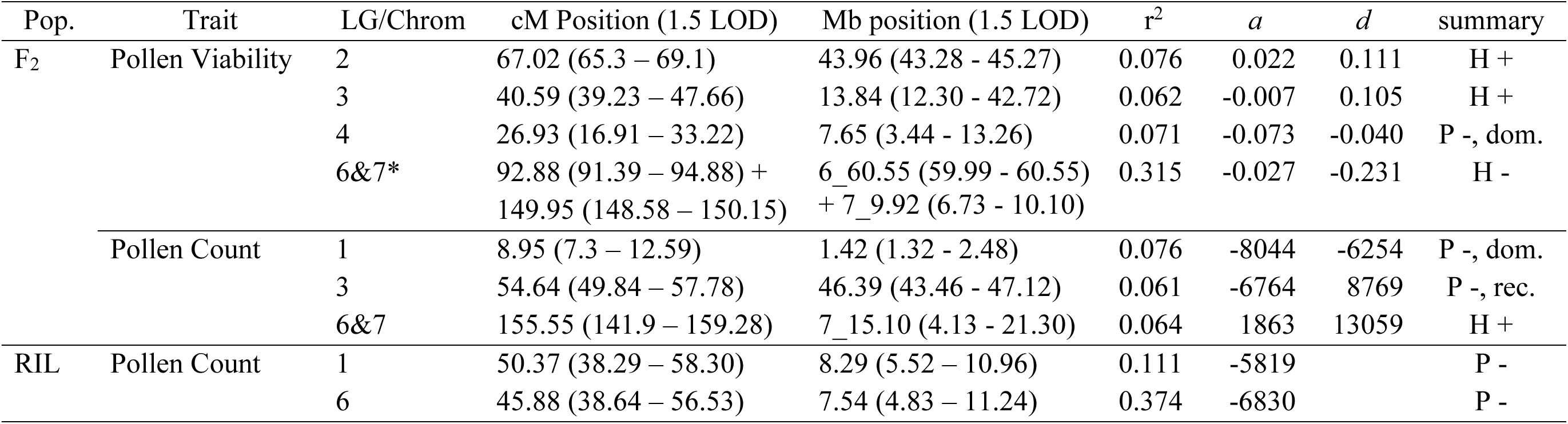
Male fertility QTL locations and effects in *M. parishii* x *M. cardinalis* UM-F_2_ and RIL populations. Additive effects (*a*) are negative when the *M. parishii* allele has the lower value, and summary indicates the overall pattern of inheritance from comparison of 2*a* vs. *a*+*d*). For the translocation-associated pollen viability QTL (6&7*) with a split location, effects are given for the higher peak. Dominance was not calculated in the RIL QTL model, which excluded heterozygotes.

The RILs were, on average, highly fertile compared to the F_2_ hybrids (mean pollen viability = 0.76; Figure S4), and we detected no significant RIL pollen viability QTLs in WinQTLCart (Figure S6). However, marker genotypes across the translocation region exhibited significant associations with pollen viability (26/42 significant at uncorrected P < 0.05, 18/42 at FDR-corrected P < 0.05). For example, at the peak marker close to the Chr7 breakpoint (Chr7_10.47), heterozygotes were ∼25% less pollen-fertile, on average, than the parental genotypes (H: 0.64± 0.04, n = 18; C = 0.79 ± 0.04, n = 18, P = 0.79 ± 0.03, n = 32). Markers on Chr6 showed parallel patterns, though they showed less distortion against *M. cardinalis* homozygotes. Although some RILs genotyped as heterozygous for LG6&7 markers were highly fertile, consistent with segregation to homozygosity between the genotyped and phenotyped generation (see Methods) and/or stabilization of compatible chromosomal configurations, the underdominant fertility effects of the translocation persist.

Consistent with the parental species’ difference in mating system, pollen number in the UM-F_2_ experiment was nearly 7-fold higher in *M. cardinalis* than in *M. parishii* (mean pollen number: CE10 = 50.38 x 10^3^, *N* = 2; PAR = 7.62 x 10^3^, *N* = 8). Pollen number in the F_2_ hybrids was intermediate (mean =38.94 x 10^3^, *N* = 245; Figure S4), and we mapped three QTLs that together explain only ∼20% of the F_2_ variance (Figure 4; Table 2), consistent with a polygenic basis to quantitative divergence in pollen production. For two of the three F_2_ QTLs, *M. parishii* homozygotes had the lowest pollen counts, while the other (on LG6&7) appeared overdominant (heterozygotes producing most pollen), and there were no significant interactions among QTLs in pairwise tests In the RIL grow-out, pollen counts were overall higher but followed the same pattern, with *M. cardinalis* ∼7x as pollen-productive and hybrids generally intermediate (mean pollen number: RILs = 19.02 x 10^3^, *N* = 100; CE10 = 32.26 x 10^3^ *N* = 17; PAR = 4.39 x 10^3^, *N*=11; Figure S4). QTLs on LG1 and LG6 together explain ∼48% of the RIL variance in pollen production, with *M. parishii* homozygotes at each producing less pollen and no evidence of interactions (Table 2; Figure S6). The genetic bases of variation in pollen number and viability were largely independent, with no significant phenotypic correlation and, with the exception of the region of suppressed recombination near translocation breakpoints, no QTL coincidence in the F_2_ hybrids (Table 2, Figure 4).

Notably, F_2_ pollen viability (or pollen number) QTLs cannot account for the LG4-LG8 gametic incompatibility. Although the LG4 pollen viability QTL is coincident with the LG4 gametic incompatibility locus at ∼6.8 Mb, individuals carrying least one *M. parishii* allele have reduced pollen viability regardless of their LG8 genotype, producing a *M. parishii*-dominant effect (Table 2; Figure S5b). Due to the lack of F_1_-recombinant P;C gametes (and P;C, H:C, and P;H F_2_s), the H;H F_2_ class is the only one capable of forming the incompatible gamete combination (i.e., 1/4 should be missing or sterile). Instead, P;P individuals (all gametes compatible) were just as male-fertile (mean pollen viability = 0.48 ± 0.03 SE, *N* = 24) as the H:H class (mean = 0.49 ± 0.02, *N* = 53), consistent with an alternative cause.

## Discussion

### *Mimulus parishii* x *M. cardinalis* genetic maps and recombinant inbred lines (RILs): a resource for understanding extreme floral divergence and speciation

The monkeyflowers of *Mimulus* Section Erythranthe have been a model system for understanding plant speciation for over five decades, with primary focus on the floral and elevational adaptation of parapatric *M. lewisii* and *M. cardinalis*. Here, we develop foundational genetic maps and RIL resources for the far more florally divergent, yet sympatric and hybridizing, pair of *M. parishii* and *M. cardinalis*. Despite structural divergence and substantial hybrid incompatibilities, we constructed consistent and high-quality ddRAD-based genetic maps to guide future investigations of the genetic and molecular mechanisms of floral, mating system, and life history evolution. Previous genetic maps in the *Erythranthe* group have used sparse PCR-based (Bradshaw *et al*. 1998; Fishman *et al*. 2013, 2015) or targeted capture markers; successful extension of this high-density gene-centric ddRAD genotyping technique from *M. guttatus* (Kolis *et al*. 2022) to the larger genomes of *M. cardinalis* and *M. parishii* holds promise for mapping and population genomics across *Mimulus*. The mapped RILs are now a permanent resource for research on adaptation and speciation, particularly the study of fitness-relevant traits that benefit from replication of recombinant genotypes under different biotic or abiotic conditions. Further, the extreme phenotypic diversity of these recombinant plants, their ease of culture, and the growing knowledge base of this flagship system make the RILs valuable for course-based undergraduate research experiences (CUREs) in genetics and evolution.

Overall, patterns of transmission ratio distortion, pseudo-linkage, and hybrid sterility reveal substantial genic and chromosomal incompatibilities between this pair of naturally hybridizing taxa. Furthermore, with the exception of the Chromosome 6-7 translocation and consistent *M. cardinalis* excess in the *YUP* supergene region of Chromosome 4 (Liang *et al*. 2023)the specific loci and patterns of hybrid sterility and TRD appeared mostly idiosyncratic to this *M. parishii* x *M. cardinalis* cross, rather than shared with either *M. lewisii* x *M. cardinalis* (Fishman *et al*. 2013) or *M. parishii* x *M. lewisii* hybrids (Fishman *et al*. 2015). Because hybrids between *M. cardinalis* and *M. lewisii* segregate for an additional translocation involving Chr 1 and Chr 8, severe underdominant pollen sterility likely overshadows other factors influencing the proportion of viable pollen (as each grain can only be sterile once) as well as patterns of marker TRD on Chromosome 8. However, we can conclude that at least two major postzygotic incompatibilities are *not* shared: the LG4-LG8 sex-independent gametophytic lethal detected here by TRD and LD (see more below) and the absent-here cytoplasmic male sterility (CMS) causing anther sterility and zero pollen production in a fraction of *M. parishii* x *M. lewisii* hybrids (Fishman *et al*. 2015). Here, the small-effect pollen number QTLs all have allelic effects consistent with *M. parishii* (also the organelle-donating grandparent, as in the *M. parishii* x *M. lewisii* CMS-producing cross) evolving lower pollen number as part of the selfing syndrome (Sicard and Lenhard 2011). In addition, although there is abundant and consistent TRD in *M. parishii* x *M cardinalis* hybrids, we did not recapitulate strong gametophytic TRD favoring *M. cardinalis* alleles on LG3 in *M. lewisii* x *M. cardinalis* hybrids (Fishman *et al*. 2013). In that cross, patterns of TRD in additional backcrosses revealed the mechanism to be single-locus and strictly male-gametophytic, suggesting a locus involved in the evolution of faster pollen tube growth in the longer-styled hummingbird pollinated species (Fishman *et al*. 2013). The absence of a parallel pattern here, where the parental difference in pollen tube growth should (if anything) be accentuated by greater divergence in flower size and mating system, suggests that the previously reported *M. cardinalis* pollen growth advantage against *M. lewisii* was dependent on the stylar or genetic context. Similarly, novel strong TRD favoring *M. parishii* homozygotes on Chr5 in our RILs (vs. the opposite pattern in F_2_ hybrids and in *M. parishii* x *M. lewisii* hybrids) suggests specific selection generated by either the greenhouse environment at UGA or the process of RIL formation. Further targeted comparisons of such barrier loci across generations, as well as among the hybrids of these three closely-related *Erythranthe* taxa, hold great promise for peeling away the complex layers of genetic interactions that shape hybrid genomes and phenotypes.

### Chromosomal translocations: a major and remarkably persistent cause of hybrid sterility

Consistent with previous coarse mapping in *M. lewisii* x *M. cardinalis* and newly available genomes (Figure 1), our results demonstrate that *M. cardinalis* carries a unique and strongly underdominant reciprocal translocation relative to closely related taxa. Inversions and translocations were among the first posited mechanisms of postzygotic reproductive isolation, as they provide cytogenetically visible evidence of hybrid genomic incompatibility (Stebbins 1958; White 1968; Grant 1971; King 1993). Although empirical studies in plant hybrids often find species-polymorphic and species-diagnostic inversions without underdominant effects on fertility (Zhang *et al*. 2021), translocations have been consistently associated with severe underdominant sterility (Lai *et al*. 2005; Fishman *et al*. 2013). Because the underdominant fertility of translocations is a direct effect of chromosomal pairing in heterozygotes (Stathos and Fishman 2014), their initial spread post-mutation should be strongly opposed by selection (Lande 1984) and remains mysterious. Notably, both hummingbird-pollinated *M. cardinalis* (Chr 6 and 7) and bee-pollinated *M. lewisii* (Chr 1 and 8) each have at least one derived reciprocal translocation, while selfer *M. parishii* carries no novel major rearrangements; thus, strong drift likely does not explain the origin of translocations in this system. Furthermore, this study shows that enforced selfing after hybridization does not eliminate this chromosomal incompatibility and its fitness effects. Thus, rather than quickly sorting to parental chromosomes, translocations may have complex and persistent effects on fertility, recombination, and introgression upon secondary contact between chromosomally-divergent plants.

The *M. cardinalis* Chromosome 6-7 translocation generates complex (non-linear) linkage relationships across much of both chromosomes in F_2_ hybrids with *M. parishii* (Figure 1), exhibits elevated heterozygosity near the breakpoints (Figure 2) and causes severe underdominant F_2_ pollen inviability (Figure 4). Substantial excess heterozygosity was retained around the translocation breakpoints (Figure 2) and, while some translocation-heterozygous RILs appear to have escaped to compatible chromosomal configurations, many of them exhibited the low pollen viability of their F_2_ ancestors. Importantly, variation among heterozygous RILs did not appear to reflect the segregation of unlinked modifiers (i.e., one or the other parental genotype substantially conferring enhanced LG6&7 heterozygote fertility by specifying alternate segregation). None of the epistatic interactions between LG6&7 and each other sterility locus, had the property of mitigating or exacerbating its underdominant pollen viability effects (Figure S5). This contrasts with previous work in *M. lewisii* x *M. cardinalis* hybrids, where *M. cardinalis* alleles on LG2 specifically accentuated the underdominant effects of the *M. cardinalis* translocation (Fishman *et al*. 2013).

This result suggests that translocations may have surprisingly persistent effects on fertility (as well as extended effects on patterns of genomic diversity and introgression) when selfing follows hybridization between structurally-divergent incipient species. Population genomics shows that *M. parishii* has recently captured a Southern Californian *M. cardinalis* chloroplast, as well as nuclear regions (Nelson *et al*. 2021b), suggesting a history of hybridization involving initial F_1_ formation with *M. cardinalis* as the seed parent, then recurrent backcrossing to *M. parishii* and/or selfing (plus possibly selection for *M. parishii*-like phenotypes). Since the CE10 chromosomal structure appears widespread throughout the *M. cardinalis* range (Fishman *et al*. 2013), we would predict that this genomic region may exhibit elevated diversity and/or unusual patterns of introgression where *M. parishii* hybridizes naturally with its hummingbird-pollinated relative. This scenario contrasts with the sorting expected for a typical barrier locus and suggests that translocations may behave differently from inversions associated with ecological barriers during speciation and secondary contact, at least in self-compatible plant species. Now that the interchange breakpoints between these naturally hybridizing taxa have been genetically and physically localized, such predictions are testable with dense sampling in areas of sympatry between *M. cardinalis* and structurally divergent species.

### A novel sex-independent gametophytic incompatibility

Patterns of TRD in both F_2_ hybrids and RILs indicate absolute non-transmission of both male and female gametes with both *M. cardinalis* alleles on Chr 8 and *M. parishii* alleles on Chr 4 (Figure 2, Tables 1 and S6). This extreme pattern of distortion is consistent with a 2-locus sex-independent gametophytic incompatibility. In land plants, which have a life cycle with two multicellular stages (a.k.a. alternation of generations), incompatibilities may act in the haploid gametophyte or in the diploid sporophyte to cause sterility. Unlike in animal sperm or egg cells (Braun *et al*. 1989), a substantial proportion of the genome is haploid-expressed in plant gametophytes (Honys and Twell 2004; Wuest *et al*. 2010; Rutley and Twell 2015). Reflecting this expression activity, most hybrid sterility loci in rice are gametophytic (Ouyang and Zhang 2013) and genome-wide patterns of TRD in multiple F_2_ populations of *Arabidopsis lyrata* also seem to be caused by gametic incompatibilities (Leppälä *et al*. 2013). Abundant gametophytic incompatibilities in plants might be due to the relatively large mutational target of haploid expressed genes, to rapid divergent evolution of gametophytic genes under efficient sex-specific selection (Gossmann *et al*. 2013, 2016) or other natural selection (Immler and Otto 2018; Beaudry *et al*. 2020), or simply to the greater opportunity for exposure of negative allelic interactions in haploid hybrid tissues (similar to hemizygosity as an explanation for Haldane’s rule).

The sex-independence of our new *M. parishii-M. cardinalis* incompatibility implies a shared molecular mechanism for loss of both male and female gamete function – but what could that be? Meiotic and developmental programs resulting in embryo sacs (female) and pollen (male) occur in distinct sexual organs of the flower, and sex-specificity of hybrid sterility loci is predominant in taxa that have been systematically studied, such as rice (Myint and Koide 2022). Nevertheless, sex-independent incompatibility loci have been identified in tomato (Rick 1966), rice (Sano 1990; Koide *et al*. 2008b), lettuce (Giesbers *et al*. 2018) and yellow monkeyflowers (Kerwin and Sweigart 2017). Because core cellular processes such as meiosis and mitosis are critical for gametogenesis in both sexes (Drews et al. 1998), many of the same genes are essential in both sexes (Drews and Yadegari 2002; Ma *et al*. 2021; Yu *et al*. 2022; Zhou *et al*. 2022). Thus, breakdown of these processes in hybrids could cause sex-independent incompatibility. However, because the LG4-LG8 gametic incompatibility does not colocalize with QTLs for either pollen number or viability, it implies a non-mitotic or meiotic mechanism that does not induce pollen abortion or substantially impair pollen development (at least in a way that could be assessed by our starch stain for viability). This contrasts with the *hms1-hms2* system in yellow monkeyflowers (Kerwin and Sweigart 2017), which causes both sex-independent TRD and pollen sterility, but parallels a sex-independent, two-locus gametophytic barrier between wild and cultivated lettuce(Giesbers *et al*. 2018). The lack of a pollen-staining effect in both this study and lettuce intriguingly suggests the involvement of genes required specifically for the functioning of mature male and female gametophytes (e.g., genes involved in both female gametogenesis and pollen tube growth/guidance; (Leszczuk *et al*. 2019; Chen *et al*. 2020). Like a lethal incompatibility causing albinism in some hybrids between yellow monkeyflowers *M. nasutus* and *M. guttatus* (Zuellig and Sweigart 2018), the completeness of gamete failure might suggest duplication and reciprocal deletion of a haploid-essential gene as a potential mechanism. While the completely non-transmitted region on Chr 8 contains ∼1000 genes flanking a putatively centromeric region of low recombination, the interacting locus on Chr 4 is ∼1 Mb containing only 80 genes. Given that parallels to other gametic lethals suggest a finite pool of functional candidate genes, the genetic and molecular mechanisms are thus amenable to further dissection.

## DATA AVAILABILITY

Raw sequence data from F_2_ and RIL mapping populations have been archived on the NCBI Sequence Read Archive under PRJNA936857 and PRJNA948041, respectively. The genotype and phenotype data for linkage, transmission ratio distortion, and QTL mapping are archived on Dryad (doi:10.5061/dryad.v6wwpzh1m) and will be made publicly available upon publication.

## ACKNOWLEDGMENTS

We thank T. C. Nelson, K. Baesen and A. Demaree for lab assistance and O. Moynahan of the UM ECOR plant growth facility, M.Moriarty, M. Opel, and C. Liu of the UConn Botanical Conservatory, and Mike Boyd and Greg Cousins of the UGA Plant Sciences Greenhouse for assistance with plant care. We are especially indebted to S. McCann for his help in generating the RILs.

---

**Figure S1.**
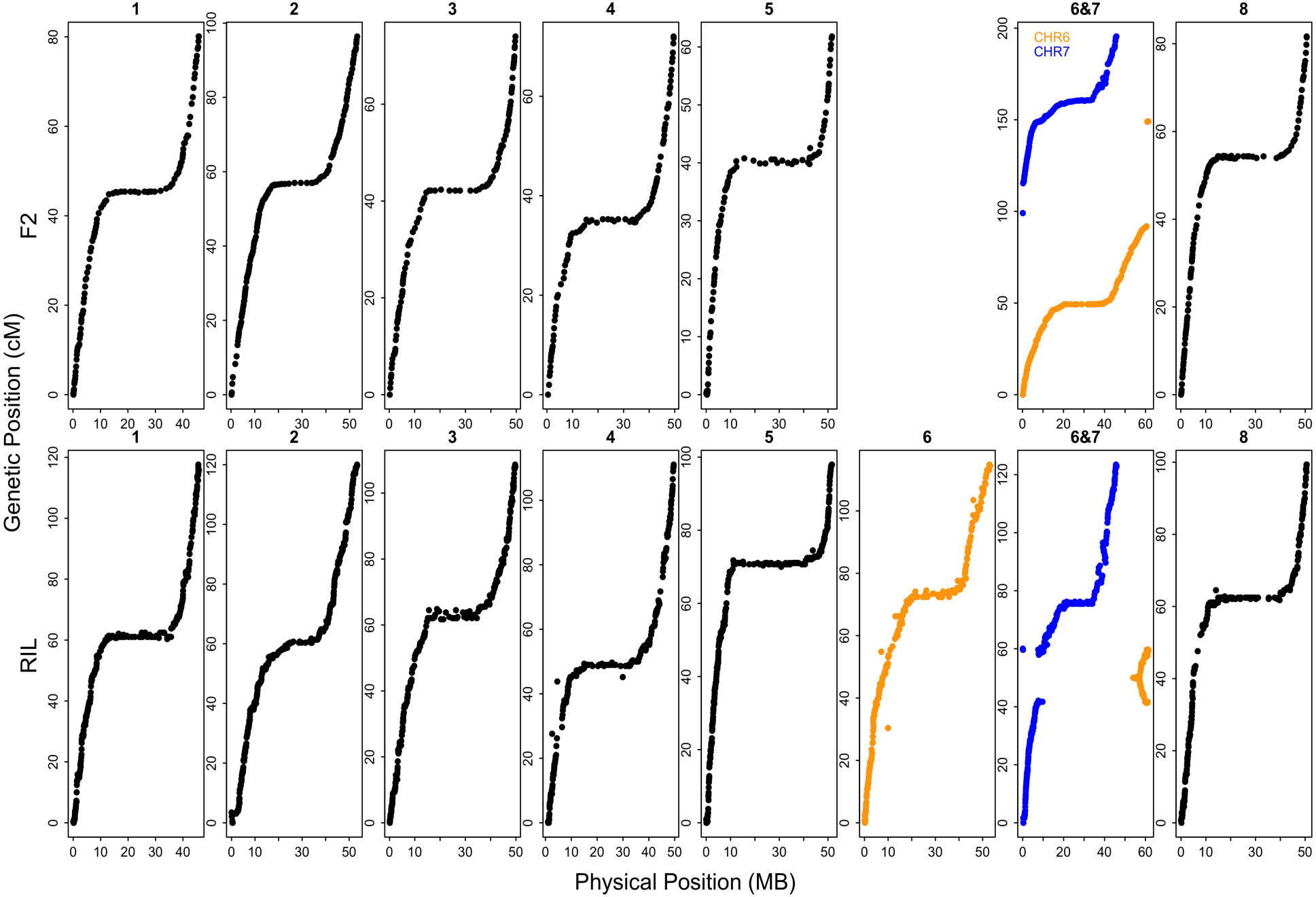
Genetic positions (cM) versus physical marker positions for F_2_ (top row) and RIL (bottom row) linkage maps, showing generally high concordance outside the translocation involving Chromosomes 6 and 7 (highlighted in color).

**Figure S2.**
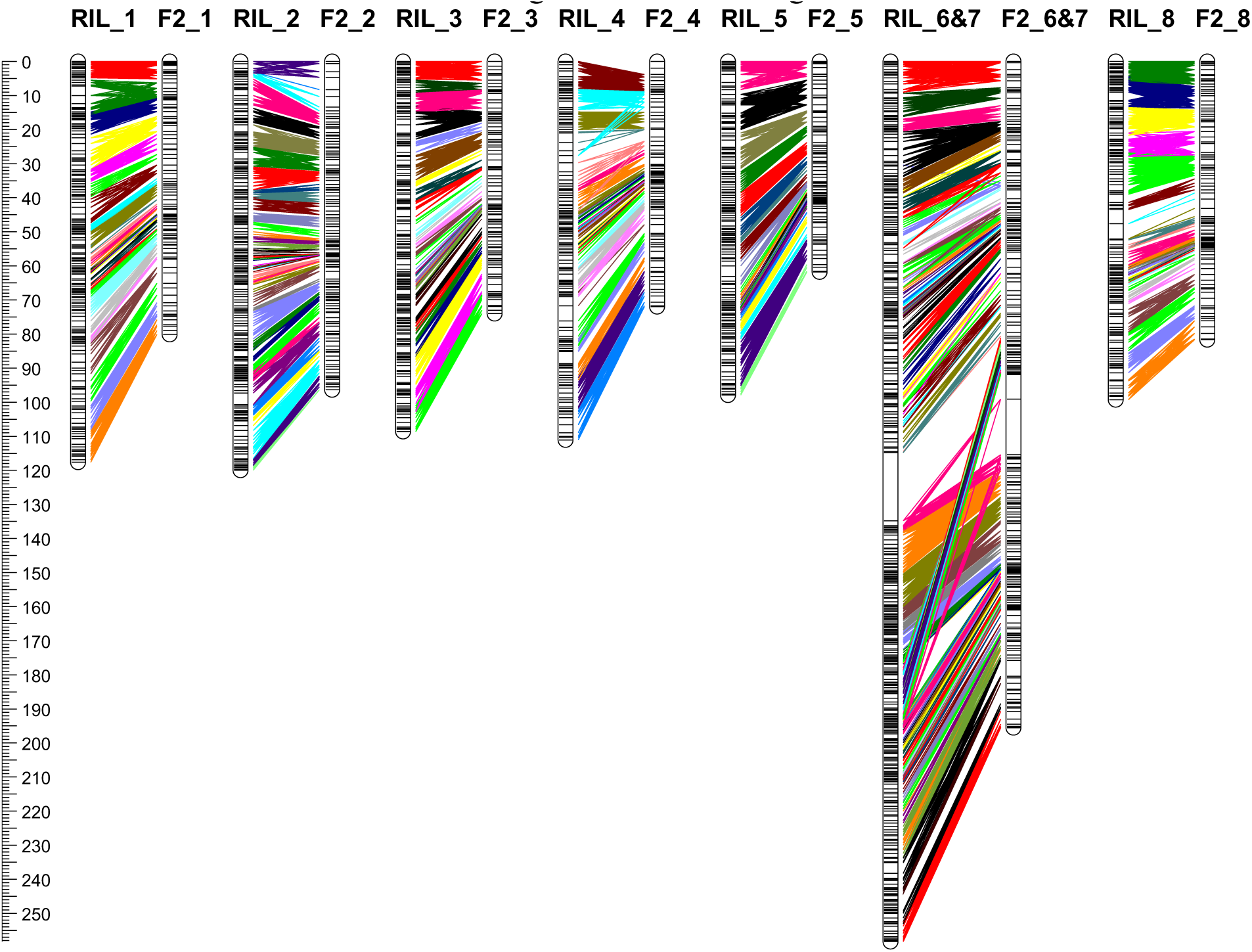
Line-up of *M. parishii* x *M. cardinalis* F_2_ and RIL linkage maps, showing their overall collinearity. Lines connect markers with shared physical positions in 1 Mb windows (arbitrarily colored for visibility) within chromosomes. A 20cM gap was inserted between concatenated RIL LG 6 and RIL LG6-7 to facilitate the alignment with the single F_2_ LG6&7.

**Figure S3.**
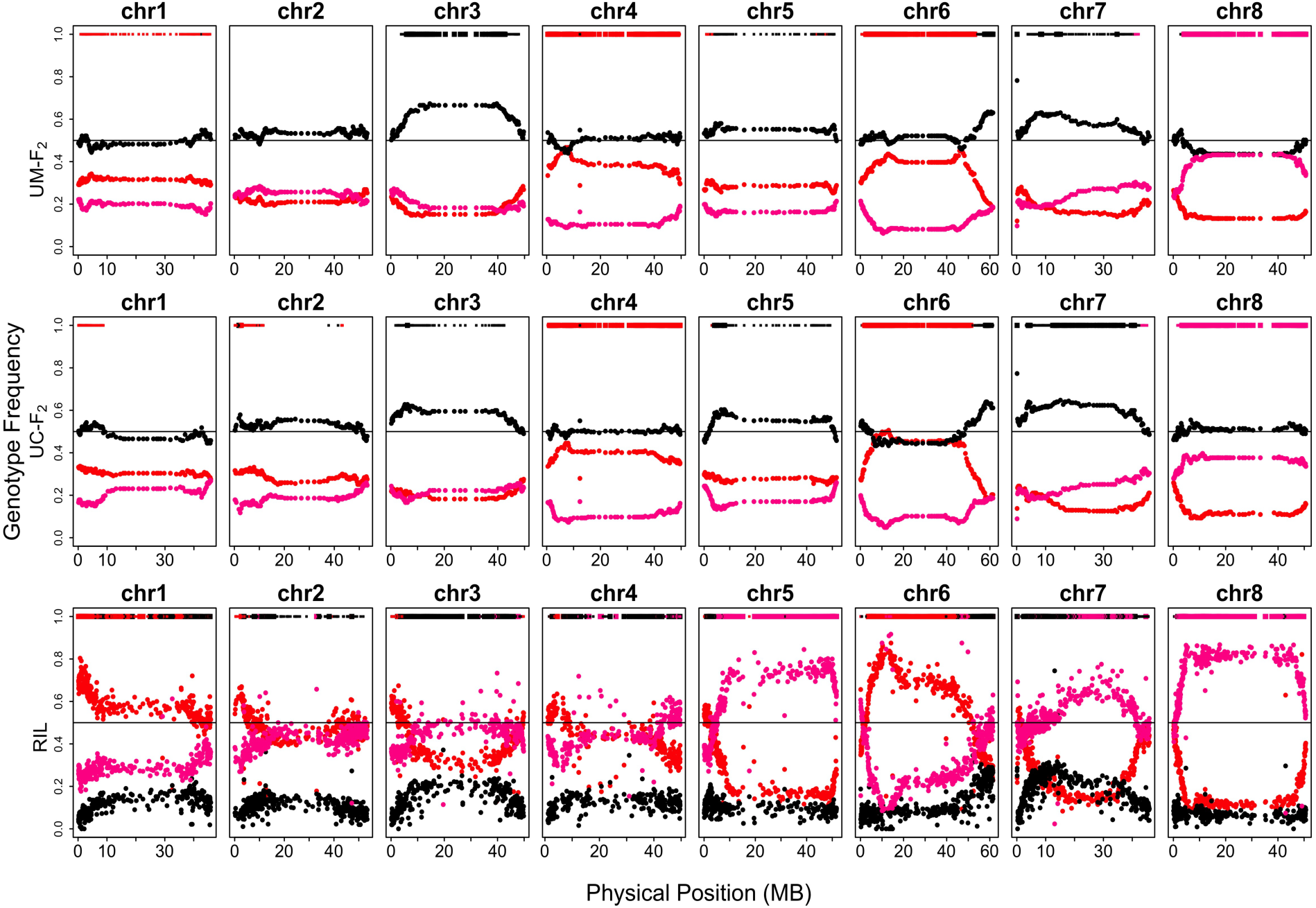
Genotype frequencies in the F_2_ and RIL populations across the genome plotted by physical position (Mb, CE10g_v2.0) across all eight chromosomes. Red dots indicate homozygous *M. cardinalis*, pink is homozygous *M. parishii*, and black is heterozygous. The bars at the top of each plot indicate regions with significant TRD by chi-squared tests at a = 0.01 (thin lines, critical value = 6.635) and at a more stringent, Bonferonni-corrected level (thick lines, F_2_ critical value = 16.442, RIL critical value = 18.217. Bar color indicates an excess of *M. cardinalis* homozygotes (red), *M. parishii* homozygotes (pink), or heterozygotes (black).

**Figure S4.**
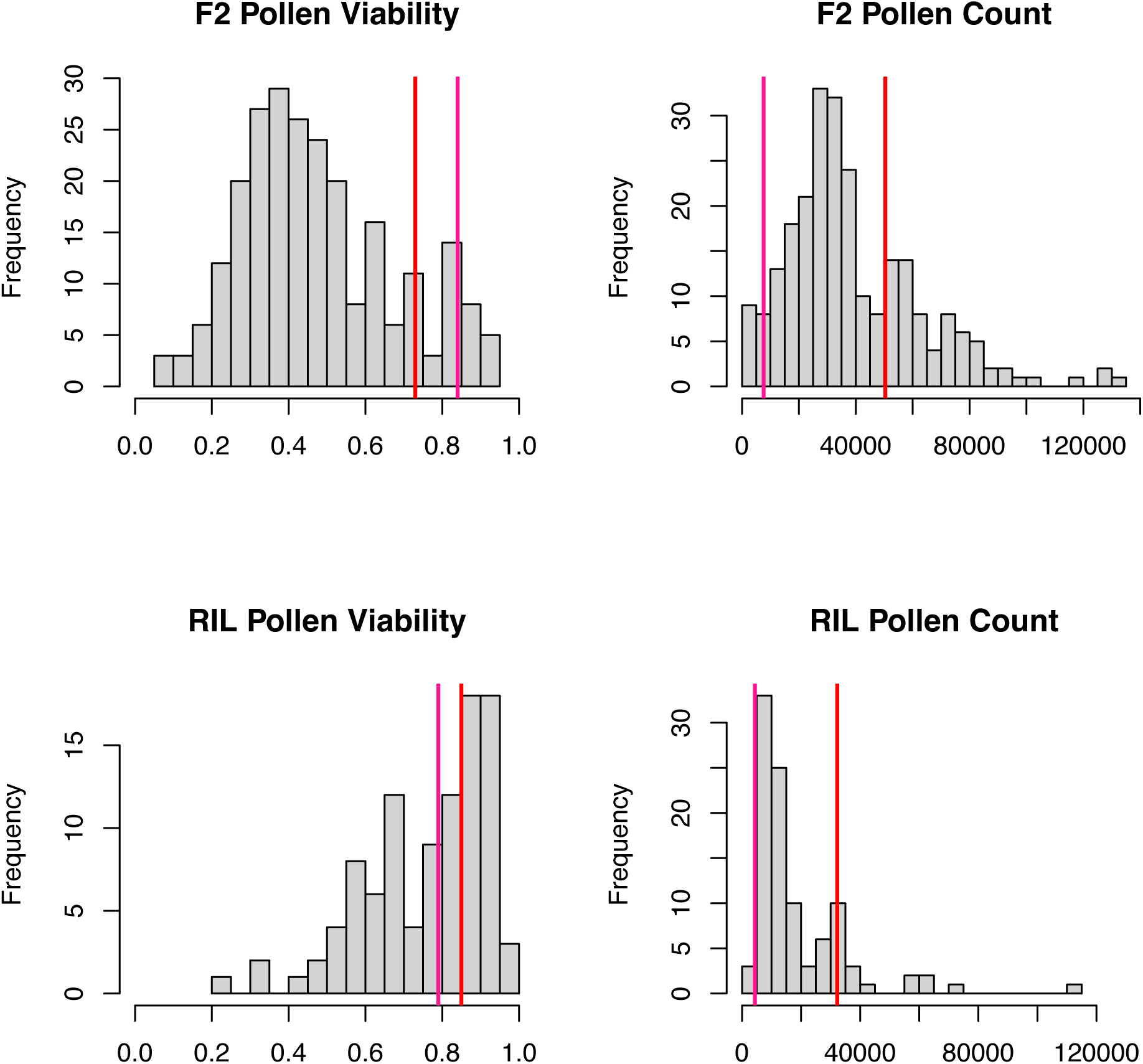
Distributions of pollen viability and pollen number for the UM-F_2_ and RIL populations. Pink lines indicate *M. parishii* average values and red lines indicate *M. cardinalis* average values.

**Figure S5.**
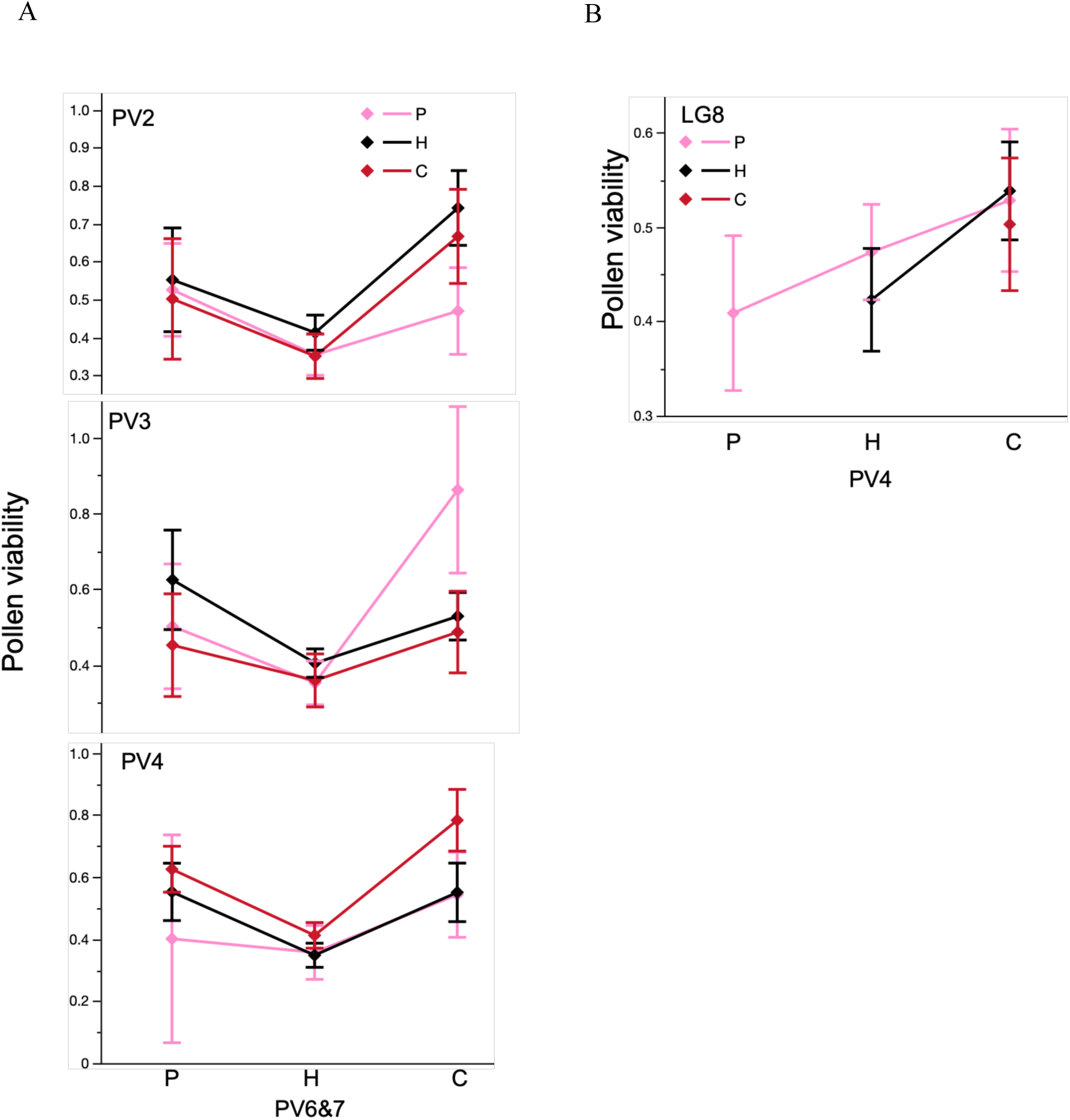
Interaction effects for pollen viability QTLs in the *M. parishii* x *M. cardinalis* UM-F_2_ mapping population. A) Significant interactions between the translocation-associated pollen inviability QTL (PV6&7) and unlinked QTLs (PV2, PV3, PV4) do not modify its underdominant effects. Graphs show least squared means and standard errors from best fit model including all four QTLs and significant interactions. B) The pollen viability QTL on LG4 (PV4) does not simply explain the LG4-LG8 gametic lethality, as that model predicts lowest fitness in the double heterozygotes rather than double *M. parishii* homozygotes. Three genotypic classes are missing due to the gametic incompatibility.

**Figure S6.**
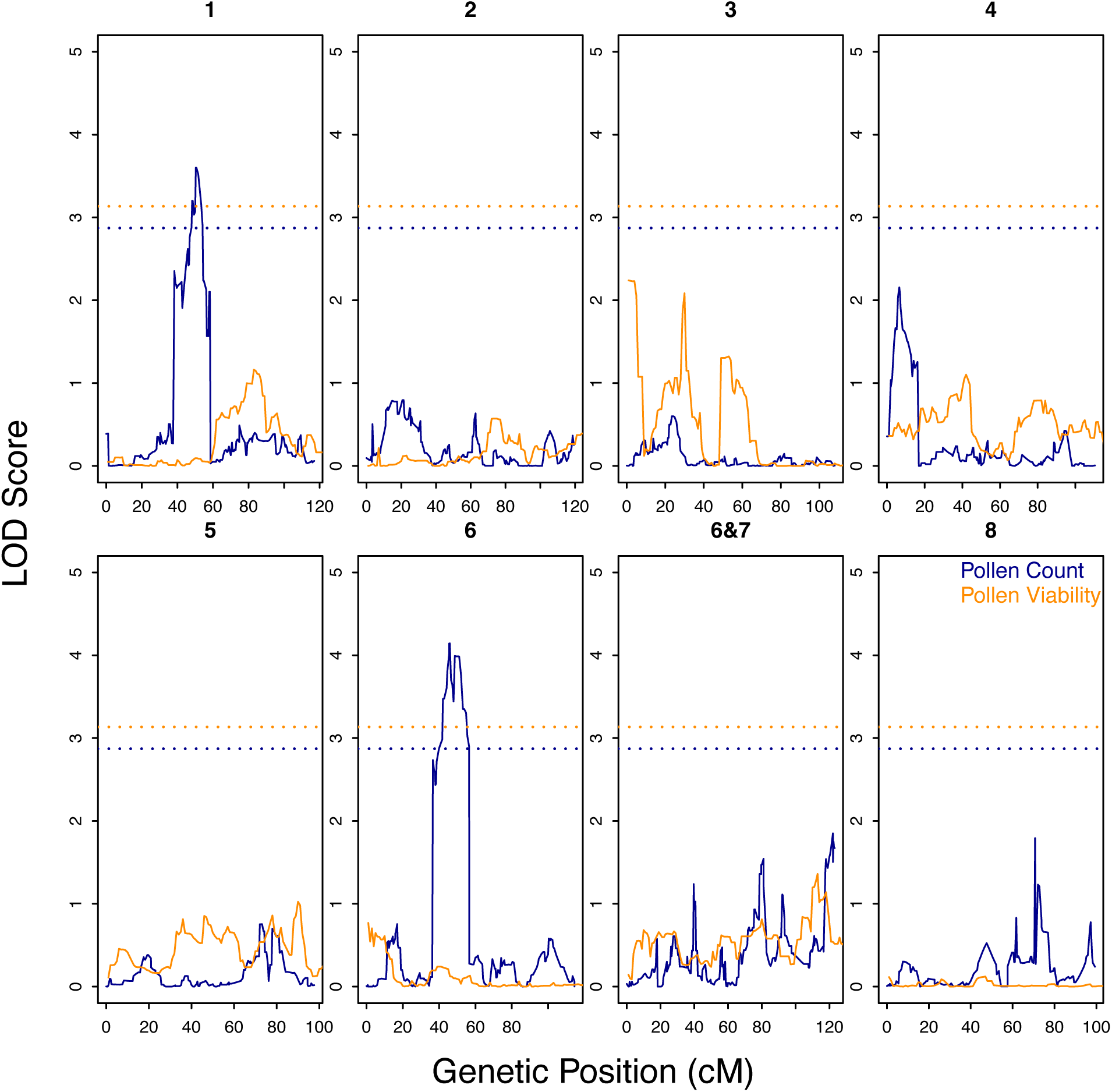
RIL LOD traces (solid lines) and significance thresholds (dotted lines) for pollen viability and pollen count QTL analyses. Orange lines are for pollen viability with the horizontal hashed line indicating the significant LOD threshold (3.13); dark blue lines are for pollen count with the horizontal hashed line indicating the significant LOD threshold (2.87).

**Table S1.** Genomic coordinates of schematic shown in Figure 1a. Coordinates are bp positions for CE10g_v2.0 and Mparg_v2.0 (mimubase.org). Regions of homology were determined by dot plot comparisons using D-Genies (Cabanettes and Klopp 2018)

| color | *M. cardinalis* (CE10) |  |  | *M. parishii* (PAR) |  |  |
| --- | --- | --- | --- | --- | --- | --- |
|  | chr | start | end | chr | start | end |
| blue | 6 | 0 | 61077219 | 6 | 0 | 58334723 |
| purple | 6 | 61515096 | 61078180 | 7 | 195334 | 625301 |
| green | 7 | 0 | 176662 | 7 | 0 | 155165 |
| red | 7 | 8434192 | 200695 | 6 | 58335309 | 65498417 |
| orange | 7 | 8436523 | 45947006 | 7 | 625481 | 34026340 |

**Table S2.** Regions (cM) in each linkage group (LG) in the F_2_ and RIL populations that show significant distortion (direction) via Chi-Square test at a = 0.01 (critical value = 6.635). Also shown are physical locations (Chrom. = chromosome number, MB = megabases).

| LG | Chrom. | cM | MB | Direction | Population |
| --- | --- | --- | --- | --- | --- |
| 1 | 1 | 4.22-77.72 | 0.87-45.56 | *M. cardinalis* | UM-F2 |
| 3 | 3 | 17.39-60.52 | 3.87-47.84 | Heterozygote | UM-F2 |
| 4 | 4 | 0-70.65 | 0.42-49.47 | *M. cardinalis* | UM-F2 |
| 5 | 5 | 3.73-24.57 | 0.96-4.3 | *M. cardinalis* | UM-F2 |
| 5 | 5 | 24.96-40.09 | 4.33-41.82 | Heterozygote | UM-F2 |
| 5 | 5 | 40.19-47.26 | 36.99-48.75 | *M. cardinalis* | UM-F2 |
| 5 | 5 | 48.63-56.79 | 49.14-50.74 | Heterozygote | UM-F2 |
| 6&7 | 6 | 3.83-81.16 | 0.62-54.26 | *M. cardinalis* | UM-F2 |
| 6&7 | 6 | 81.85-91.87 | 54.58-60.55 | Heterozygote | UM-F2 |
| 6&7 | 7 | 99.09 | 0.151 | Heterozygote | UM-F2 |
| 6&7 | 7 | 140.61-180.9 | 3.85-42.07 | Heterozygote | UM-F2 |
| 8 | 8 | 19.16-81.65 | 2.81-50.56 | *M. parishii* | UM-F2 |
| 1 | 1 | 0-39.2 | 0.18-8.71 | *M. cardinalis* | UC-F2 |
| 2 | 2 | 0-47.66 | 0.23-11.6 | *M. cardinalis* | UC-F2 |
| 2 | 2 | 10.324 | 2.35 | Heterozygote | UC-F2 |
| 2 | 2 | 58.466 | 37.74 | Heterozygote | UC-F2 |
| 2 | 2 | 61.61-65.25 | 41.51-43.28 | *M. cardinalis* | UC-F2 |
| 3 | 3 | 8.25-47.06 | 1.81-42.38 | Heterozygote | UC-F2 |
| 4 | 4 | 0-72.02 | 0.42-49.56 | *M. cardinalis* | UC-F2 |
| 5 | 5 | 17.69-21.62 | 2.98-3.66 | *M. cardinalis* | UC-F2 |
| 5 | 5 | 23.78-48.63 | 4.07-49.14 | Heterozygote | UC-F2 |
| 6&7 | 6 | 4.62-79.49 | 0.72-52.96 | *M. cardinalis* | UC-F2 |
| 6&7 | 6 | 80.57-91.87 | 53.68-60.55 | Heterozygote | UC-F2 |
| 6&7 | 7 | 99.09 | 0.15 | Heterozygote | UC-F2 |
| 6&7 | 7 | 135.6-182.48 | 3.25-42.65 | Heterozygote | UC-F2 |
| 6&7 | 7 | 184.15-189.75 | 43.09-44.98 | *M. parishii* | UC-F2 |
| 8 | 8 | 11.59-81.65 | 1.62-50.56 | *M. parishii* | UC-F2 |
| 1 | 1 | 0 - 44.838 | 0.156732 - 6.553678 | *M. cardinalis* | RIL |
| 1 | 1 | 47.942 - 79.325 | 6.856646 - 40.837789 | *M. cardinalis* | RIL |
| 1 | 1 | 81.049 - 91.742 | 41.709833 - 43.096019 | *M. cardinalis* | RIL |
| 1 | 1 | 103.124 - 106.229 | 44.436729 - 44.665482 | *M. cardinalis* | RIL |
| 1 | 1 | 107.608 - 107.953 | 44.758167 - 44.739669 | Heterozygote | RIL |
| 1 | 1 | 116.92 - 117.61 | 45.56426 - 45.582184 | *M. cardinalis* | RIL |
| 2 | 2 | 0 - 0.69 | 0.373876 - 0.373816 | *M. cardinalis* | RIL |
| 2 | 2 | 2.759 - 8.277 | 0.77023 - 3.393731 | *M. cardinalis* | RIL |
| 2 | 2 | 11.036 - 12.415 | 3.491784 - 3.954329 | *M. parishii* | RIL |
| 2 | 2 | 16.899 - 17.588 | 4.834793 - 5.004551 | *M. cardinalis* | RIL |
| 2 | 2 | 25.176 - 25.521 | 6.002301 - 6.295216 | *M. cardinalis* | RIL |
| 2 | 2 | 45.868 - 47.937 | 11.32465 - 12.738106 | Heterozygote | RIL |
| 2 | 2 | 51.386 - 51.731 | 13.777678 - 13.468346 | Heterozygote | RIL |
| 2 | 2 | 90.699 - 91.044 | 46.767261 - 47.190381 | *M. cardinalis* | RIL |
| 3 | 3 | 0 - 0.69 | 0.11538 - 0.20371 | *M. cardinalis* | RIL |
| 3 | 3 | 1.725 - 5.173 | 0.405792 - 0.995216 | *M. cardinalis* | RIL |
| 3 | 3 | 6.208 - 8.622 | 1.114955 - 1.356512 | *M. cardinalis* | RIL |
| 3 | 3 | 13.105 - 13.449 | 2.81225 - 2.792177 | Heterozygote | RIL |
| 3 | 3 | 18.623 - 22.073 | 3.252869 - 3.665279 | *M. parishii* | RIL |
| 3 | 3 | 26.556 - 27.591 | 4.880219 - 5.041717 | *M. cardinalis* | RIL |
| 3 | 3 | 35.868 - 36.213 | 6.49436 - 6.795835 | Heterozygote | RIL |
| 3 | 3 | 37.592 - 37.937 | 6.996131 - 7.148653 | *M. cardinalis* | RIL |
| 3 | 3 | 38.972 - 43.455 | 7.243214 - 8.836678 | Heterozygote | RIL |
| 3 | 3 | 46.56 - 50.699 | 9.000542 - 10.257358 | Heterozygote | RIL |
| 3 | 3 | 52.078 - 66.908 | 10.864504 - 39.47233 | Heterozygote | RIL |
| 3 | 3 | 68.287 - 73.46 | 40.46393 - 42.321623 | Heterozygote | RIL |
| 3 | 3 | 75.184 - 76.909 | 42.979279 - 44.142058 | Heterozygote | RIL |
| 4 | 4 | 0 - 0.345 | 1.296198 - 1.258052 | *M. cardinalis* | RIL |
| 4 | 4 | 13.45 - 15.519 | 2.854257 - 3.190946 | *M. cardinalis* | RIL |
| 4 | 4 | 17.933 - 20.692 | 3.439869 - 4.126026 | *M. cardinalis* | RIL |
| 4 | 4 | 23.798 - 35.874 | 4.225103 - 6.971637 | *M. cardinalis* | RIL |
| 4 | 4 | 37.253 - 37.598 | 7.977272 - 8.154489 | Heterozygote | RIL |
| 4 | 4 | 44.151 - 45.875 | 9.197184 - 11.797988 | Heterozygote | RIL |
| 4 | 4 | 47.254 - 47.254 | 12.037848 - 12.689921 | *M. parishii* | RIL |
| 4 | 4 | 48.633 - 50.013 | 18.206963 - 35.752834 | *M. parishii* | RIL |
| 4 | 4 | 55.185 - 56.22 | 40.174398 - 39.960666 | Heterozygote | RIL |
| 4 | 4 | 64.842 - 65.877 | 42.745021 - 42.939379 | Heterozygote | RIL |
| 4 | 4 | 77.961 - 78.996 | 45.426916 - 45.646088 | Heterozygote | RIL |
| 4 | 4 | 81.411 - 83.135 | 46.103086 - 45.959864 | *M. parishii* | RIL |
| 4 | 4 | 84.859 - 85.204 | 46.757086 - 46.838726 | *M. parishii* | RIL |
| 4 | 4 | 91.072 - 98.659 | 47.607254 - 48.699141 | *M. parishii* | RIL |
| 4 | 4 | 101.418 - 101.763 | 48.918801 - 49.040036 | Heterozygote | RIL |
| 4 | 4 | 106.592 - 111.077 | 49.233258 - 49.620274 | *M. parishii* | RIL |
| 5 | 5 | 0.345 - 0.69 | 0.261829 - 0.353901 | *M. cardinalis* | RIL |
| 5 | 5 | 6.209 - 7.588 | 0.954502 - 0.954494 | *M. cardinalis* | RIL |
| 5 | 5 | 16.212 - 18.281 | 1.379473 - 1.685151 | Heterozygote | RIL |
| 5 | 5 | 19.66 - 20.005 | 1.743674 - 1.863926 | Heterozygote | RIL |
| 5 | 5 | 21.04 - 21.04 | 2.023362 - 2.076503 | Heterozygote | RIL |
| 5 | 5 | 22.074 - 25.869 | 2.337737 - 2.488783 | Heterozygote | RIL |
| 5 | 5 | 28.283 - 28.972 | 2.777281 - 2.839236 | Heterozygote | RIL |
| 5 | 5 | 49.319 - 97.948 | 5.321919 - 51.659755 | *M. parishii* | RIL |
| 6 | 6 | 0 - 0.69 | 0.239398 - 0.092231 | *M. parishii* | RIL |
| 6 | 6 | 1.379 - 2.414 | 0.332759 - 0.107857 | *M. parishii* | RIL |
| 6 | 6 | 25.521 - 114.847 | 3.412154 - 52.901735 | *M. cardinalis* | RIL |
| 8 | 8 | 5.863 - 7.587 | 1.337734 - 1.52118 | *M. parishii* | RIL |
| 8 | 8 | 11.382 - 95.898 | 1.637985 - 50.276892 | Heterozygote | RIL |
| 6&7 | 7 | 21.381 - 22.761 | 2.660981 - 2.780974 | Heterozygote | RIL |
| 6&7 | 6 and 7 | 24.485 - 75.523 | 2.897287 - 24.606753 | Heterozygote | RIL |
| 6&7 | 7 | 75.523 - 98.629 | 24.606759 - 40.984997 | *M. parishii* | RIL |

**Table S3.** Regions (cM) in each linkage group (LG) in the F2 and RIL populations that show significant distortion (direction) via Chi-Square test at the Bonferonni-corrected critical values (F_2_ critical value = 16.442, RIL critical value = 18.217). Also shown are physical locations (Chrom. = chromosome number, MB = megabases).

| LG | Chrom. | cM | MB | Direction | Population |
| --- | --- | --- | --- | --- | --- |
| 3 | 3 | 25.05-47.65 | 5.86-42.72 | Heterozygote | UM-F2 |
| 4 | 4 | 0-65.34 | 0.418-48.8 | *M. cardinalis* | UM-F2 |
| 6&7 | 6 | 15.43-79.49 | 2.52-52.96 | *M. cardinalis* | UM-F2 |
| 6&7 | 6 | 88.33-91.87 | 57.54-60.55 | Heterozygote | UM-F2 |
| 6&7 | 7 | 99.09 | 0.1511 | Heterozygote | UM-F2 |
| 6&7 | 6 and 7 | 149.06 | (6) 60.98 - (7) 8.63 | Heterozygote | UM-F2 |
| 6&7 | 7 | 154.07-155.54 | 13.8-15.1 | Heterozygote | UM-F2 |
| 6&7 | 7 | 162.61 | 34.64 | Heterozygote | UM-F2 |
| 8 | 8 | 27.81-81.46 | 4.04-50.63 | *M. parishii* | UM-F2 |
| 2 | 2 | 8.26-13.37 | 1.82-2.77 | Heterozygote | UC-F2 |
| 3 | 3 | 26.13 | 6.25 | Heterozygote | UC-F2 |
| 4 | 4 | 3.83-72.02 | 1.04-49.56 | *M. cardinalis* | UC-F2 |
| 5 | 5 | 23.78-35.47 | 4.07-8.01 | Heterozygote | UC-F2 |
| 6&7 | 6 | 12.97-73.79 | 2.06-51.51 | *M. cardinalis* | UC-F2 |
| 6&7 | 6 | 89.22-91.87 | 57.92-60.55 | Heterozygote | UC-F2 |
| 6&7 | 7 | 99.09 | 0.15 | Heterozygote | UC-F2 |
| 6&7 | 7 | 141.01-143.85 | 39.77-46.67 | Heterozygote | UC-F2 |
| 6&7 | 7 | 152.49-166.45 | 12.43-36.69 | Heterozygote | UC-F2 |
| 6&7 | 7 | 170.18-172.34 | 40.43-40.65 | Heterozygote | UC-F2 |
| 8 | 8 | 22.2-79.79 | 3.28-50.46 | *M. parishii* | UC-F2 |
| 1 | 1 | 0 - 26.214 | 0.156732 - 3.25118 | *M. cardinalis* | RIL |
| 1 | 1 | 61.047 - 61.047 | 15.839755 - 35.671627 | Heterozygote | RIL |
| 1 | 1 | 64.151 - 64.151 | 36.017146 - 36.260853 | Heterozygote | RIL |
| 1 | 1 | 65.185 - 65.53 | 37.107993 - 37.107849 | Heterozygote | RIL |
| 1 | 1 | 67.254 - 68.289 | 38.057857 - 38.857561 | Heterozygote | RIL |
| 1 | 1 | 70.703 - 71.047 | 39.350951 - 39.350949 | *M. cardinalis* | RIL |
| 1 | 1 | 76.22 - 77.945 | 40.020711 - 40.180308 | Heterozygote | RIL |
| 1 | 1 | 28.284 - 28.973 | 3.255494 - 3.801921 | *M. cardinalis* | RIL |
| 1 | 1 | 31.733 - 32.422 | 3.959125 - 4.254112 | *M. cardinalis* | RIL |
| 1 | 1 | 41.388 - 42.768 | 6.312352 - 6.467571 | *M. cardinalis* | RIL |
| 2 | 2 | 3.104 - 3.794 | 1.483501 - 2.767148 | *M. cardinalis* | RIL |
| 2 | 2 | 47.247 - 47.247 | 12.42425 - 12.490094 | Heterozygote | RIL |
| 3 | 3 | 68.632 - 68.977 | 40.575854 - 40.463884 | Heterozygote | RIL |
| 3 | 3 | 41.041 - 41.386 | 7.46192 - 8.159341 | Heterozygote | RIL |
| 3 | 3 | 54.837 - 54.837 | 12.482446 - 13.121749 | Heterozygote | RIL |
| 3 | 3 | 57.251 - 59.666 | 13.985791 - 14.809159 | Heterozygote | RIL |
| 3 | 3 | 61.735 - 66.908 | 21.314691 - 39.341831 | Heterozygote | RIL |
| 4 | 4 | 48.633 - 48.978 | 20.327014 - 23.925429 | Heterozygote | RIL |
| 5 | 5 | 54.836 - 54.836 | 7.071539 - 7.417066 | *M. parishii* | RIL |
| 5 | 5 | 60.009 - 96.914 | 8.307822 - 51.266631 | *M. parishii* | RIL |
| 6 | 6 | 34.488 - 53.801 | 4.113997 - 10.303714 | *M. cardinalis* | RIL |
| 6 | 6 | 96.223 - 97.602 | 45.66901 - 47.040811 | *M. cardinalis* | RIL |
| 6 | 6 | 111.743 - 114.847 | 51.575546 - 52.901735 | Heterozygote | RIL |
| 6 | 6 | 58.975 - 62.079 | 12.866981 - 14.881438 | *M. cardinalis* | RIL |
| 6 | 6 | 63.804 - 64.838 | 15.807607 - 15.805921 | *M. cardinalis* | RIL |
| 6 | 6 | 66.563 - 76.218 | 16.230404 - 41.90439 | *M. cardinalis* | RIL |
| 6 | 6 | 77.598 - 77.943 | 41.153674 - 42.26249 | *M. cardinalis* | RIL |
| 6 | 6 | 79.667 - 88.635 | 42.509669 - 44.418568 | *M. cardinalis* | RIL |
| 8 | 8 | 18.624 - 33.108 | 2.751408 - 4.532002 | *M. parishii* | RIL |
| 8 | 8 | 35.177 - 90.034 | 4.532018 - 49.890287 | *M. parishii* | RIL |
| 6&7 | 7 | 25.519 - 27.589 | 3.1644 - 3.536587 | Heterozygote | RIL |
| 6&7 | 7 | 28.623 - 30.692 | 3.907913 - 4.406577 | Heterozygote | RIL |
| 6&7 | 6 and 7 | 31.037 - 75.523 | 4.168792 - 29.659926 | Heterozygote | RIL |
| 6&7 | 7 | 75.523 - 88.628 | 29.659956 - 38.014756 | *M. parishii* | RIL |

**Table S4.** Average interchromosomal LD (*r*) across all pairwise marker comparisons for the UM-F2 population for those which are greater than the 95% quantile (0.176). ‘N’ is the total number of pairwise marker comparisons and ‘N_95q’ is the number of those pairwise comparisons that are greater than the 95% quantile. ‘NA’ indicates there were no pairwise marker comparisons greater than the 95% quantile.

| LG Comparison | LD ($r$) | N | N_95q | Proportion |
| --- | --- | --- | --- | --- |
| 1_2 | 0.1957 | 15594 | 341 | 0.0219 |
| 1_3 | NA | 13108 | NA | NA |
| 1_4 | 0.1997 | 11526 | 65 | 0.0056 |
| 1_5 | 0.1782 | 12091 | 1 | 0.0001 |
| 1_6&7 | 0.1866 | 32883 | 61 | 0.0019 |
| 1_8 | NA | 14690 | NA | NA |
| 2_3 | 0.1868 | 16008 | 17 | 0.0011 |
| 2_4 | 0.1826 | 14076 | 5 | 0.0004 |
| 2_5 | 0.1861 | 14766 | 15 | 0.001 |
| 2_6&7 | 0.1821 | 40158 | 329 | 0.0082 |
| 2_8 | 0.2068 | 17940 | 1423 | 0.0793 |
| 3_4 | 0.1857 | 11832 | 247 | 0.0209 |
| 3_5 | NA | 12412 | NA | NA |
| 3_6&7 | 0.1802 | 33756 | 16 | 0.0005 |
| 3_8 | 0.1785 | 15080 | 6 | 0.0004 |
| 4_5 | NA | 10914 | NA | NA |
| 4_6&7 | 0.186 | 29682 | 68 | 0.0023 |
| 4_8 | 0.2788 | 13260 | 7805 | 0.5886 |
| 5_6&7 | 0.1944 | 31137 | 69 | 0.0022 |
| 5_8 | 0.1824 | 13910 | 78 | 0.0056 |
| 6&7_8 | 0.1809 | 37830 | 6 | 0.0002 |

**Table S5.** Average interchromosomal LD (*r*) across all pairwise marker comparisons for the RIL population for those which are greater than the 95% quantile (0.305). ‘N’ is the total number of pairwise marker comparisons and ‘N_95q’ is the number of those pairwise comparisons that are greater than the 95% quantile. ‘NA’ indicates there were no pairwise marker comparisons greater than the 95% quantile.

| LG<br>Comparison | LD ($r$) | N | N_95q | Proportion |
| --- | --- | --- | --- | --- |
| 1_2 | 0.3273 | 79 | 102315 | 0.0008 |
| 1_3 | 0.4133 | 223 | 84930 | 0.0026 |
| 1_4 | 0.4697 | 277 | 67830 | 0.0041 |
| 1_5 | 0.3593 | 150 | 80370 | 0.0019 |
| 1_6 | 0.3308 | 540 | 80085 | 0.0067 |
| 1_6&7 | 0.3393 | 232 | 140220 | 0.0017 |
| 1_8 | 0.4116 | 119 | 85500 | 0.0014 |
| 2_3 | 0.3826 | 28 | 106982 | 0.0003 |
| 2_4 | 0.3572 | 3387 | 85442 | 0.0396 |
| 2_5 | 0.4327 | 1104 | 101238 | 0.0109 |
| 2_6 | 0.3734 | 2621 | 100879 | 0.026 |
| 2_6&7 | 0.3617 | 1241 | 176628 | 0.007 |
| 2_8 | 0.3282 | 637 | 107700 | 0.0059 |
| 3_4 | 0.3837 | 418 | 70924 | 0.0059 |
| 3_5 | 0.3943 | 972 | 84036 | 0.0116 |
| 3_6 | 0.3572 | 220 | 83738 | 0.0026 |
| 3_6&7 | 0.3474 | 13747 | 146616 | 0.0938 |
| 3_8 | 0.3552 | 1820 | 89400 | 0.0204 |
| 4_5 | 0.3712 | 15126 | 67116 | 0.2254 |
| 4_6 | 0.3314 | 2004 | 66878 | 0.03 |
| 4_6&7 | 0.3511 | 9731 | 117096 | 0.0831 |
| 4_8 | 0.3532 | 12411 | 71400 | 0.1738 |
| 5_6 | 0.3885 | 771 | 79242 | 0.0097 |
| 5_6&7 | 0.3623 | 10785 | 138744 | 0.0777 |
| 5_8 | 0.3474 | 5668 | 84600 | 0.067 |
| 6_6&7 | 0.35 | 3592 | 138252 | 0.026 |
| 6_8 | 0.46 | 1008 | 84300 | 0.012 |
| 6&7_8 | 0.3571 | 24246 | 147600 | 0.1643 |

**Table S6.** Two-locus genotype (C is *M. cardinalis*, P is *M. parishii*, and H is heterozygote) counts and frequencies (in parentheses) for each of the nine possible genotype combinations between linkage groups 4 and 8 for the UM-F_2_ and RIL populations. Data shown are the observed counts and frequencies, expected counts and frequencies under normal Mendelian inheritance, expected counts and frequencies under a non-Mendelian inheritance model of gametic inviability (P;C gametes inviable), and expected counts and frequencies under a non-Mendelian inheritance model of zygotic inviability (P_;C_ gametes inviable).

| Genotype<br>(LG4; LG8) | F<sub>2</sub> Population |  |  |  | RIL Population |  |  |  |
| --- | --- | --- | --- | --- | --- | --- | --- | --- |
|  | Observed | Mendelian | P;C gametes<br>inviable | P_<sub>;</sub>C_<sub>;</sub><br>zygotes<br>inviable | Observed | Mendelian | P;C gametes<br>inviable | P_<sub>;</sub>C_<sub>;</sub><br>zygotes<br>inviable |
| C;C | 24 (0.09) | 16.06<br>(0.0625) | 28.56 (0.11) | 36.71 (0.143) | 18 (0.13) | 31.29 (0.23) | 41.72 (0.30) | 41.72 (0.30) |
| H;C | 0 (0) | 32.13 (0.125) | 0 (0) | 0 (0) | 0 (0) | 2.14 (0.015) | 0 (0) | 0 (0) |
| P;C | 0 (0) | 16.06<br>(0.0625) | 0 (0) | 0 (0) | 0 (0) | 31.29 (0.23) | 0 (0) | 0 (0) |
| C;H | 61 (0.24) | 32.13 (0.125) | 57.11 (0.22) | 73.43 (0.286) | 6 (0.04) | 2.14 (0.015) | 4.28 (0.031) | 6.42 (0.046) |
| H;H | 53 (0.21) | 64.25 (0.25) | 57.11 (0.22) | 0 (0) | 6 (0.04) | 4.28 (0.031) | 4.28 (0.031) | 0 (0) |
| P;H | 0 (0) | 32.13 (0.125) | 0 (0) | 0 (0) | 0 (0) | 2.14 (0.015) | 0 (0) | 0 (0) |
| C;P | 27 (0.11) | 16.06<br>(0.0625) | 28.56 (0.11) | 36.71 (0.143) | 40 (0.29) | 31.29 (0.23) | 41.72 (0.30) | 41.72 (0.30) |
| H;P | 58 (0.23) | 32.13 (0.125) | 57.11 (0.22) | 73.43 (0.286) | 21 (0.15) | 2.14 (0.015) | 4.28 (0.031) | 6.42 (0.046) |
| P;P | 34 (0.13) | 16.06<br>(0.0625) | 28.56 (0.11) | 36.71 (0.143) | 47 (0.34) | 31.29 (0.23) | 41.72 (0.30) | 41.72 (0.30) |

